# Personality Variation in Associative Learning and Memory Retention in Zebrafish (*Danio rerio*)

**DOI:** 10.64898/2026.09.08.750049

**Authors:** Princess Sunday-Jimmy, Tran Thien An Le, Ryan Y. Wong

**Affiliations:** University of Nebraska at Omaha Biology Department, Omaha, NE USA

**Keywords:** Associative learning, memory, personality, sex, food reward

## Abstract

Animals differ in how they acquire and retain information, and modify their behavior while interacting with their environment. These differences have been linked to personality type, yet how personality type and sex influence reward-associated memory is not well understood. We investigated the effects of personality type, sex, days of conditioning, and food reward on learning and memory retention in zebrafish using the conditioned place preference paradigm. We conditioned zebrafish to associate a visual background with a food reward across eleven sessions and assessed memory recall seven days later by measuring duration, frequency of entry, and latency to first entry in the conditioned zone. We observed that bold fish had early conditioned zone engagement, whereas shy zebrafish gradually increased their behavior into the conditioned zone with repeated conditioning. With food reward, preference for the conditioned zone increased after 7 and 11 days post-conditioning relative to baseline. Seven days after the last conditioning with food reward, the preference for conditioned zone remained unchanged in shy fish. This suggests that shy fish retain the ability to recall the learned association for at least 7 days after conditioning. Overall, personality, food reward, and sex influence behavioral changes in food reward-associated cognition, with shy-treated males showing gradual increases in behavior into the conditioned zone. These findings suggest that personality and sex differences may depend on the nature and valence of the associative task to recall. Thus, this study emphasizes the importance of assessing memory retention alongside learning acquisition to understand the relationship between personality types and cognition.

## 1. Introduction

Animals consistently face new experiences while interacting with their environment, and the ability to learn from and recall these experiences over time is important for adaptation and survival[1]. Learning acquisition and memory retention are related, but they represent distinct functional processes of cognition. An individual may quickly acquire an association but fail to retain it over time or learn slowly while still consolidating the information into a durable memory [2,3]. These dynamic patterns of learning and memory can also vary between individuals [4,5].

One major source of variation in cognition is personality type [6,7], which can shape how animals respond to stress [8–10], novelty [11–13], and reward [14]. Studies across animal taxa have linked individual variation in cognitive performance to personality type [5,15,16], particularly differences along the bold-shy continuum [6,7,11,16–18]. Bold individuals are generally less neophobic, risk-tolerant, show less stress-related and flexible behaviors. Bold individuals are also more exploratory, which may result in faster learning and decision-making, while shy individuals display the opposite behavioral traits [6,13,19–22]. These personality-related differences in cognition are often explained through a risk-reward trade-off framework, in which bold individuals may gain early access to resources through quick exploration, whereas shy individuals may reduce risk through more cautious engagement with novel stimuli [17,23–25]. Additionally, these differences in behavioral tendencies may reflect underlying variation in neurobiological systems involved in reward processing, motivation, gene expression, and stress regulation, all of which could be tied to how animals acquire and retain learned associations [26–29]. Studies in fish and birds have reported stronger recall in shy than bold individuals in memory tasks involving both aversive (conditioned fear and predatory) and positively valenced stimuli, such as food reward tasks [7,30–32]. Sex may also influence memory retention [33], although studies have reported inconsistent findings [6]. For example, a study in zebrafish found no difference in memory capacity between males and females [34] whereas another found males to exhibit higher memory retention [30]. Therefore, the extent to which personality type and sex influence memory retention is not well understood.

Zebrafish (*Danio rerio*) offer a valuable model for studying the interaction between personality and cognition [3,35]. Zebrafish exhibit diverse cognitive abilities, including appetitive associative learning [36], object-recognition memory [37], and episodic-like memory [38]. Consistent individual differences in boldness, behavioral flexibility, exploration, and risk-taking have been documented across selectively bred laboratory and wild zebrafish populations, supporting personality variation along the bold-shy continuum [6,39–41]. Selectively bred bold and shy zebrafish lines have been shown to differ across various biological processes, including neuromolecular response to anxiolytic drugs [42], stress and anxiety [9,43], antioxidant response [44], transcriptome profiles [28,45], hormonal responses [43], associative reward learning [6], and contextual fear learning [7]. Together, these established cognitive abilities and consistent personality differences make zebrafish well-suited for examining how personality influences associative learning and memory.

This study aimed to investigate how personality, food reward conditioning, and sex influence associative learning and memory using a conditioned place paradigm. We compared bold and shy individuals across repeated learning and memory probes using time spent in the conditioned zone, frequency of entry, and latency to first as behavioral measures of cognitive performance. We hypothesized that (i) bold individuals would display faster learning acquisition and lower memory retention due to their more exploratory behavior compared to the shy individuals, and (ii) personality, sex, and treatment would interact to influence learning and memory retention across days of conditioning.

## 2. Methods

### 2.1 Subjects

This study used zebrafish lines that were selectively bred for 13 generations in the lab from wild-caught zebrafish originating near the village of Gaighata, India (approximately 60 km from Kolkata), to exhibit on average a shy or bold personality type [46]. We randomly selected 24 fish per line for this study. Our final sample sizes consisted of 10 control bold (4 females, 6 males),10 treated bold (4 females, 6 males), 9 control shy (3 females, 6 males), and 10 treated shy (3 females, 7 males). We determined sex through visual inspection of secondary sexual characteristics [6,28]. The fish were housed together in 40 L tanks and were fed twice a day with Tetramin Tropical Flakes (Tetra, USA). One week prior to testing, we transferred each fish into an individual 3-liter tank on a recirculating water system (Pentair Aquatic Eco-Systems or Aquaneering) using UV and solid filtration on a 14:10 L/D cycle, with lights on between 06:00–20:00, temperature between 26-27 °C, and water quality of pH:6.8-7.4, conductivity: 600-900 μS. To increase feeding motivation and reduce the chances of satiation during our conditioned training, we fasted the fish four days before the start of testing.

### 2.2 Behavioral Assay (Conditioned Place Preference Task)

We assessed associative learning and memory following a previously established conditioned place preference paradigm [6]. In brief, we tested the fish individually in a Zantiks LT unit (Zantiks, Cambridge, UK). We first habituated the fish to the testing tank for 10 minutes on two consecutive days without any training light stimuli or food reward. We determined baseline preference for a visual stimulus (grey or checkered pattern) the following day by presenting both stimuli zone simultaneously for ten minutes to each fish. For each fish we assigned the conditioned (CZ) stimulus zone as the visual stimulus where the fish spent the least time during baseline. The non-conditioned stimulus zone (NZ) was assigned as the visual stimulus in which the fish spent the most amount of time.

We conducted conditioning across 11 sessions divided into three phases (3 sessions before probe 1, 4 sessions before probe 2, and 4 sessions before Probe 3). Each phase was separated by a two-day rest period during which we neither fed nor tested the fish. During each conditioning session, we exposed each fish to NZ for 5 minutes, immediately followed by the CZ for 5 minutes. Throughout the CZ 100 ul of homogenized brine shrimp solution (11.4 g frozen brine shrimp dissolved in 35 mL distilled water) was dispensed every 60 seconds as a food reward for the treatment fish. An equivalent volume of distilled water was dispensed to the control fish. We did not dispense food or distilled water during the five minutes NZ period. Control fish were fed an equal amount of brine shrimp after the conditioning session.

We conducted three probe trials (mirroring the baseline procedure) the day after the third, seventh, and eleventh conditioning sessions to assess learning acquisition. Seven days after the final conditioning session, we ran an additional probe trial (probe 4) to evaluate memory retention. Fish were not fed during the seven-day interval between the final conditioning session and probe 4, to be consistent with the fasting procedure for non-testing days. For baseline and probe trials we quantified time spent, frequency of entry, and latency to first entry into the conditioned zone using commercial software (EthoVision XT Noldus Version 17). All procedures were approved by the University of Nebraska at Omaha’s Institutional Animal Care and Use Committee.

### 2.3. Statistical Analysis

All statistical analyses were performed using R (4.5.0)[47] software in RStudio (v-2026.01.0+392)[48]. We visualized the behavioral parameters for the final 5 minutes using the ggplot2 [49] and the ggpubr packages[50] and the ggboxplot function in R. For baseline and probe trials, we analyzed only the final 5 minutes of the behavioral data to reduce the influence of handling on fish behavior. Time spent in the conditioned zone, frequency of entry into the conditioned zone, and latency to first entry into the conditioned zone were not normally distributed. Therefore, we applied a log transformation on all variables and verified they were normally distributed using the Shapiro-Wilk test. We performed a type III repeated measures ANOVA to evaluate the main and interaction effects of personality, sex, treatment, and number of conditioning sessions on time spent in, frequency of entry into, and latency to first entry into the conditioned zone. We found a main effect for sex only for frequency of entry. Therefore, sex was removed from the model after showing no significant main effect (p > 0.05) for time spent in and latency to first enter into the conditioned zone. We subsequently reported significant main and interactive effects for each behavioral measure. To investigate the direction of significant effects across and within each variable, we performed a post-hoc pairwise comparison using Tukey adjustments for multiple comparisons.

## 3. Results

### 3.1. Effects of Personality, Treatment and Conditioned Days on Time Spent in the Conditioned Zone (CZ)

We performed a three-way repeated measures ANOVA to evaluate the effects of personality, treatment, and conditioned day on time spent in the conditioned zone. There was a significant main effect of conditioned day (conditioned day: *F*_2.91,96.14_ = 4.314, p = 0.007, *n*^2^*G* = 0.094) . The post hoc test revealed a trend for increased time spent in the CZ at day 7 post-conditioning compared to baseline (p = 0.088), and a significant increase at day 11 post-conditioning (p = 0.003) compared to baseline. There was also a significant main effect of personality type (personality: *F*_1,33_=8.522, p = 0.006, *n*^2^*G* = 0.05), where bold fish spent significantly more time in the conditioned zone (p = 0.006) than the shy fish. There was no significant main effect of treatment ( *F*_1,33_ = 2.361, p = 0.134, *n*^2^*G* = 0.014) (Fig.1).

**Figure 1:**
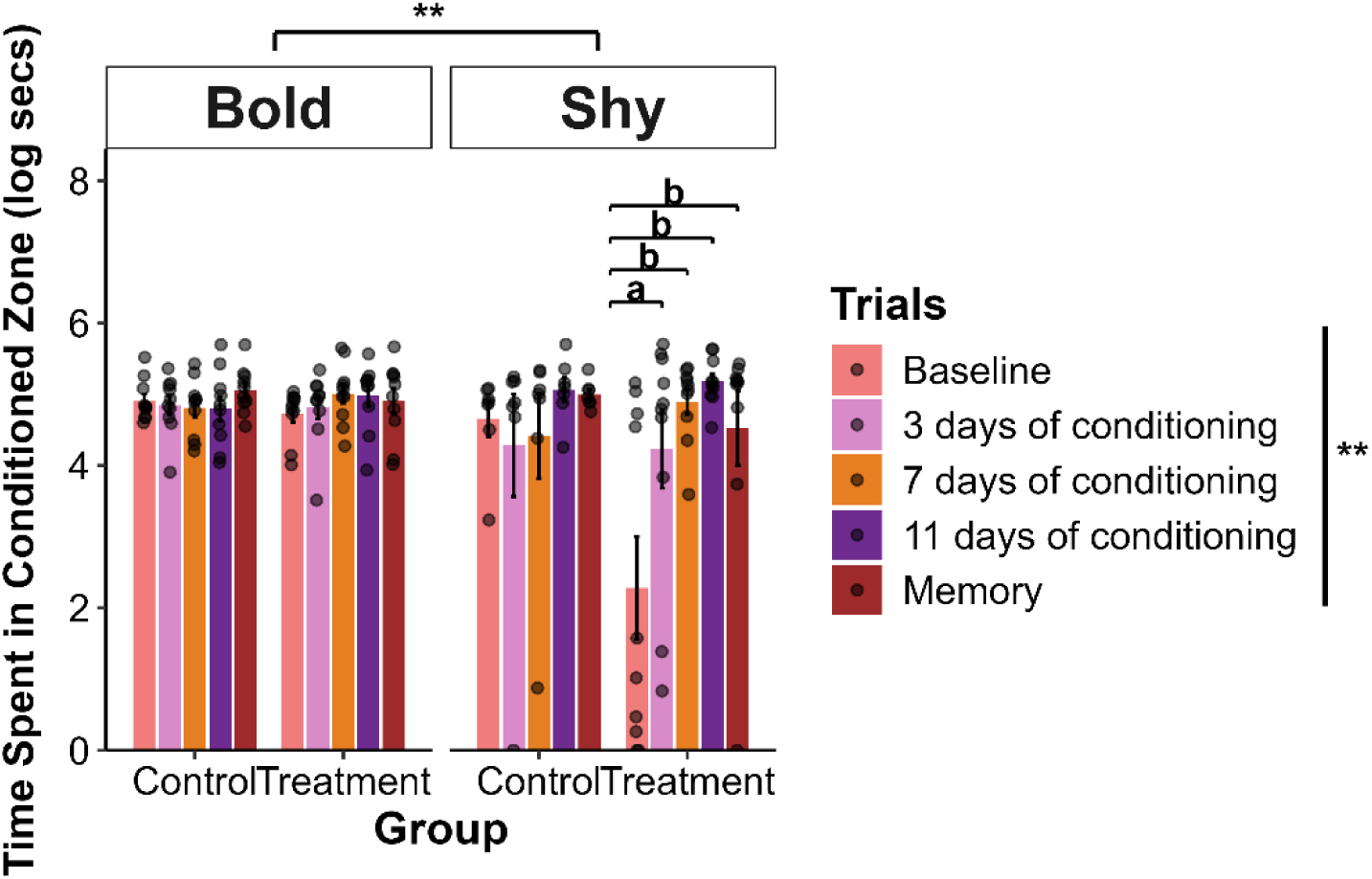
Time spent in the conditioned zone across personality, treatment, and days of conditioning. Peach bars are at baseline, pink bars represent learning after 3 days of conditioning sessions (Probe 1), orange bars are after 7 days of conditioning sessions (Probe 2), and purple bars are after 11 days of conditioning sessions (Probe 3). Red bars indicate memory retention seven days after the final conditioning session. Asterisks represent main effects of personality and conditioned days. Bars represent mean time spent in the conditioned zone during the final 5 min of each trial for bold and shy zebrafish in the control and treatment groups at each conditioned days. Error bars indicate SEM, and individual points represent individual fish.

There was a significant treatment by conditioned day interaction effect ( *F*_2.91,96.14_=3.767, p = 0.014, *n*^2^*G* = 0.083), where fish within the treatment group significantly increased time spent in the conditioned zone after 3 (p =0.05), 7 (p <0.001), and 11 (p < 0.0001) days of conditioning from baseline. Control group showed no significant difference in time spent in the conditioned zone between baseline and 3 (p = 0.982), 7 (p = 0.990), or 11 (p = 0.990) days after conditioning. For the memory trial (probe 4), there was no significant difference between the control and treatment groups (p = 0.329) in time spent in conditioned zone. Within the treatment group, there was only a significant increase in time spent in the CZ (p = 0.03) in the memory trial relative to baseline. Within the control group we did not observe any differences between baseline and probes in time spent in the conditioned zone. See supplementary table S1 for full post-hoc comparisons.

We also found significant personality type by conditioned day interaction effect on time spent in the conditioned zone ( *F*_2.91,96.14_=3.220, p = 0.027, *n*^2^*G* = 0.072). Bold fish spent more time in the conditioned zone only at baseline (p = 0.003), relative to the shy. Within the shy group, there was a significant increase in time spent in the conditioned zone after 7 (p = 0.019) and 11 (p <0.001) days of conditioning relative to baseline. For the memory trial, within the shy group, there was a significant increase in time spent in the CZ (p = 0.04) compared to the baseline.

Within the bold group we did not observe any differences between baseline and probes in time spent in the conditioned zone. See supplementary table S1 for full post-hoc comparisons.

There was a trend for significant three-way interaction effect (treatment*personality* conditioned day: *F*_2.91,96.14_= 2.259, p = 0.088, *n*^2^*G* = 0.052) on time spent in the conditioned zone. Only the shy-treated group significantly increased time in the CZ after 3 (p <0.001), 7 (p <0.0001), 11 (p < 0.0001) days of conditioning relative to baseline. Within the shy control, bold control and treated groups we did not observe any differences between baseline and probes in time spent in the conditioned zone. See supplementary table S1 for full post-hoc comparisons. For the memory trial, only the shy-treated group showed a significant increase in time spent in the CZ zone (p < 0.0001) relative to baseline (Fig.1).

### 3.2. Effects of Personality and Sex, Treatment and Conditioned Days on Entry Frequency in the Conditioned Zone (CZ)

A four-way repeated measures ANOVA was performed to evaluate the effects of personality, treatment, sex, and conditioned day on the frequency of entries into the conditioned zone. There was a significant main effect of personality ( *F*_1,29_ = 12.90, p = 0.001, *n*^2^*G* = 0.117), where the bold fish (p = 0.001) entered the CZ more than the shy fish (Fig.2). There was also a main effect of sex (sex: *F*_1,29_= 8.84, p = 0.006, *n*^2^*G* = 0.083), where the female fish (p = 0.005) significantly entered CZ more than the male fish (Fig. 2). There was no significant main effect of treatment (treatment: *F*_1,29_ = 0.486, p = 0.491, *n*^2^*G* = 0.005), or conditioned day (conditioned day: *F*_3.33,96.59_ =1.249, p=0.296, *n*^2^*G* = 0.029).

**Figure 2:**
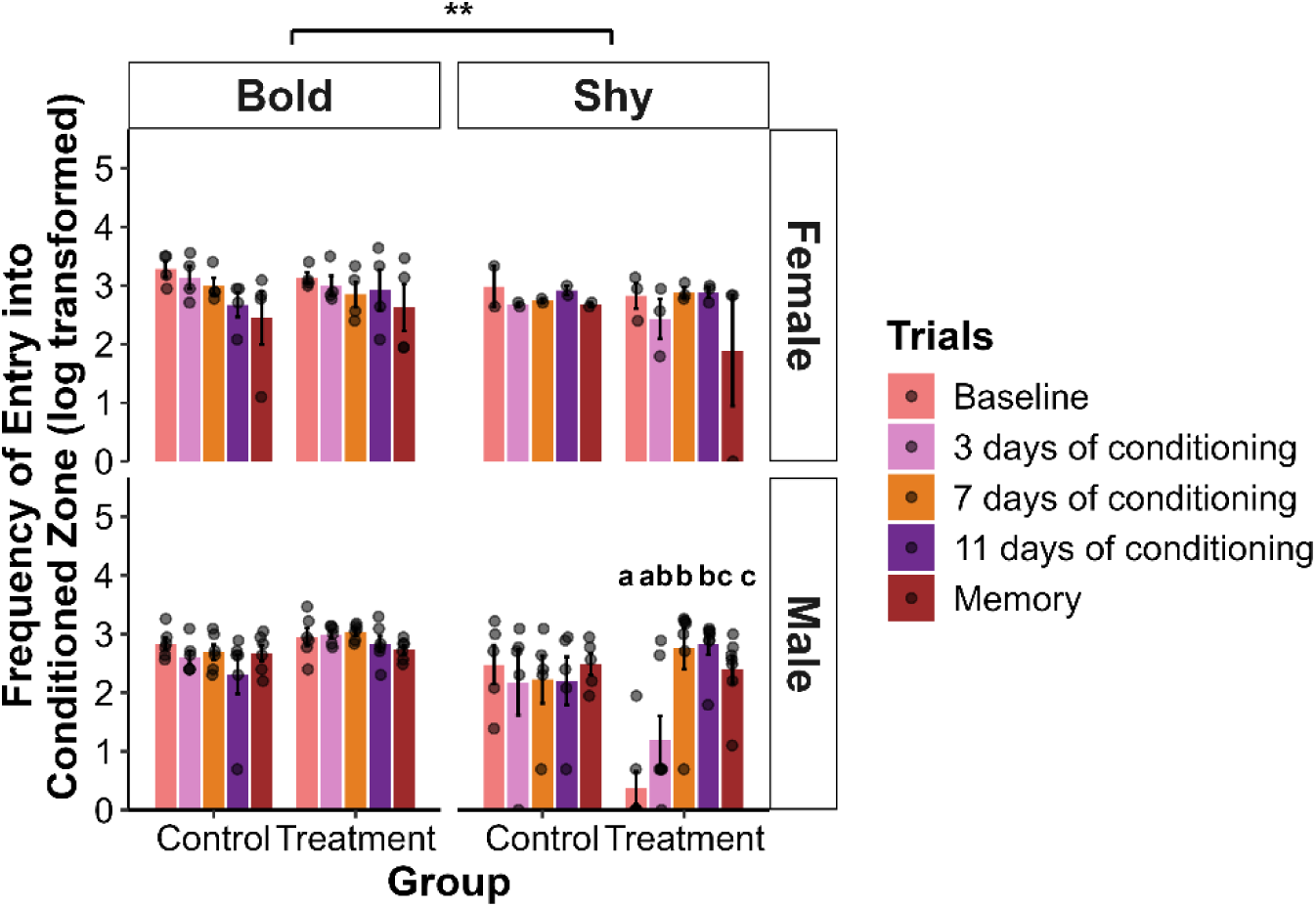
Frequency of entry into the conditioned zone across personality, sex, treatment, and conditioned day. Peach bars are at baseline, pink bars represent learning after 3 days of conditioning sessions (Probe 1), orange bars are after 7 days of conditioning sessions (Probe 2), and purple bars are after 11 days of conditioning sessions (Probe 3). Red bars indicate memory retention seven days after the final conditioning session. Bars represent mean entry frequency in the conditioned zone during the final 5 min of each trial for bold and shy zebrafish in the control and treatment groups at each conditioned days. Error bars indicate SEM, and individual points represent individual fish. Different letters indicate significant post hoc differences within shy-treated males. Asterisks indicate significant main effect of personality.

There were significant treatment by conditioned day interaction effect ( *F*_3.33,96.59_=3.139, p = 0.025, *n*^2^*G* = 0.071) on frequency of entries into the conditioned zone. The control fish had a higher frequency of entry into the conditioned zone only at baseline (p = 0.003), relative to day 3 (p = 0.355), day 7 (p =0.303), and day 11 (p = 0.122) post-conditioning. For the memory trial (probe 4), there was no significant difference between the control and treatment groups (p = 0.506) in frequency of entry into the conditioned zone. Fish within the treatment group significantly increased number of entries after 7 (p = 0.022) and 11 (p = 0.026) days of conditioning relative to baseline. Within the control group we did not observe any differences between baseline and probes in time spent in the conditioned zone. See supplementary table S2 for full post-hoc comparisons. For the memory trial, only within the treatment group, there was a trend for decreased entry frequency into the CZ compared to day 7 post-conditioning (p = 0.067). There were no significant differences when compared to baseline (p = 0.991), day 3 (p = 1.000), and day 11 post-conditioning (p = 0.202). Within the control group we did not observe any differences between baseline and probes in number of entries into the conditioned zone. See supplementary table S2 for full post-hoc comparisons.

We also found a significant personality by conditioned day interaction effect ( *F*_3.33,96.59_ = 3.604, p = 0.013, *n*^2^*G* = 0.080), where only at baseline (p < 0.0001) and after 3 days of conditioning (p = 0.004) the bold fish entered CZ significantly more than the shy fish. Within the shy group, there was a trend for increased frequency of entry after 11 days of conditioning from baseline (p = 0.007). For the memory trial, there was no significant difference in number of entries into the conditioned zone between bold and shy (p = 0.292). Within the bold group, we did not observe any differences between baseline and probes in the number of entries into the conditioned zone. See supplementary table S2 for full post-hoc comparisons.

There was a significant sex by conditioned day interaction effect ( *F*_3.33,96.59_ = 3.680, p = 0.012, *n*^2^*G* = 0.082) on number of entries into CZ. Females entered the CZ more than males at baseline (p < 0.001) and after 3 days of conditioning (p = 0.04). Within the male group, the frequency of entry increased significantly from baseline at day 7 post-conditioning (p = 0.01). Female group showed no significant difference in time spent in the conditioned zone between baseline and 3 (p = 0.875), 7(p = 0.889), or 11(p = 0.852) days after conditioning. For the memory trial, there was no significant difference between the sexes in number of entries into the CZ (p = 0.5069). Within the female group, there was a trend of decreased entry frequency into the CZ during the memory trial when compared to baseline (p = 0.008). There were no significant differences when compared to day 3 (p = 0.677), day 7 (p = 0.207), and day 11 post-conditioning (p = 0.425). Within the male group, there was a significant increase in the number of entries into the CZ 7 days after conditioning (p = 0.013) relative to baseline.

There was a trend for treatment by personality interaction effect ( *F*_1,29_ = 3.490, p = 0.072, *n*^2^*G* = 0.035), where the bold-treated (p = 0.003) entered the CZ more than shy-treated. We did not see any significant difference within the control group (p = 0.257). We also found a trend for personality by sex interaction effects ( *F*_1,29_=3.325, p = 0.079, *n*^2^*G* = 0.033), where bold males (p < 0.0001) to significantly entered the CZ more than the shy males. We did not observe any significant difference between bold and shy for the female group (p = 0.287). Only within the shy group the female increased number of entries into the CZ than males (p = 0.004). We did not observe any significant differences between sex in the bold group (p = 0.379).

There was a trend for four-way interaction effect (treatment*personality*sex*conditioned day: *F*_3.33,96.59_=2.227, p = 0.083, *n*^2^*G* = 0.051), where only the shy-treated males significantly entered the CZ more at day 7 (p<0.0001), 11 post-conditioning (p <0.0001) compared to baseline. Within the shy control, bold control and treated groups, we did not observe any differences between baseline and probes in the number of entries into the conditioned zone. See supplementary table S2 for full post-hoc comparisons. For the memory trial, only the shy-treated males significantly entered the CZ more compared to baseline (p <0.001) and when compared to day 3 post-conditioning (p = 0.04) (Fig. 2). Within the shy control sexes, bold control and treated sex, and shy treated female groups we did not observe any differences between baseline and probes in number of entries into the conditioned zone during memory trial. See supplementary table S2 for full post-hoc comparisons.

### 3.3. Effects of Personality, Treatment and Conditioned Days on Latency to first enter the Conditioned Zone (CZ)

A three-way repeated measures ANOVA was performed to evaluate the effects of personality, treatment, and conditioned day on latency to first entry in the CZ. There was a significant main effect of personality (personality: *F*_1,33_=6.411, p = 0.016, *n*^2^*G* = 0.037), where the shy fish displayed a significantly shorter latency to first enter the CZ (p = 0.016) than the bold fish (Fig.3). There was no main effects of treatment ( *F*_1,33_ = 0.137, p = 0.714, *n*^2^*G* = 0.001), conditioning days ( *F*_3.69,121.69_ = 1.640, p = 0.173, *n*^2^*G* = 0.038), or three-way interaction (personality*treatment*conditioned day: *F*_3.69,121.69_ = 1.380, p = 0.247, *n*^2^*G* = 0.032) (Fig.3).

**Figure 3:**
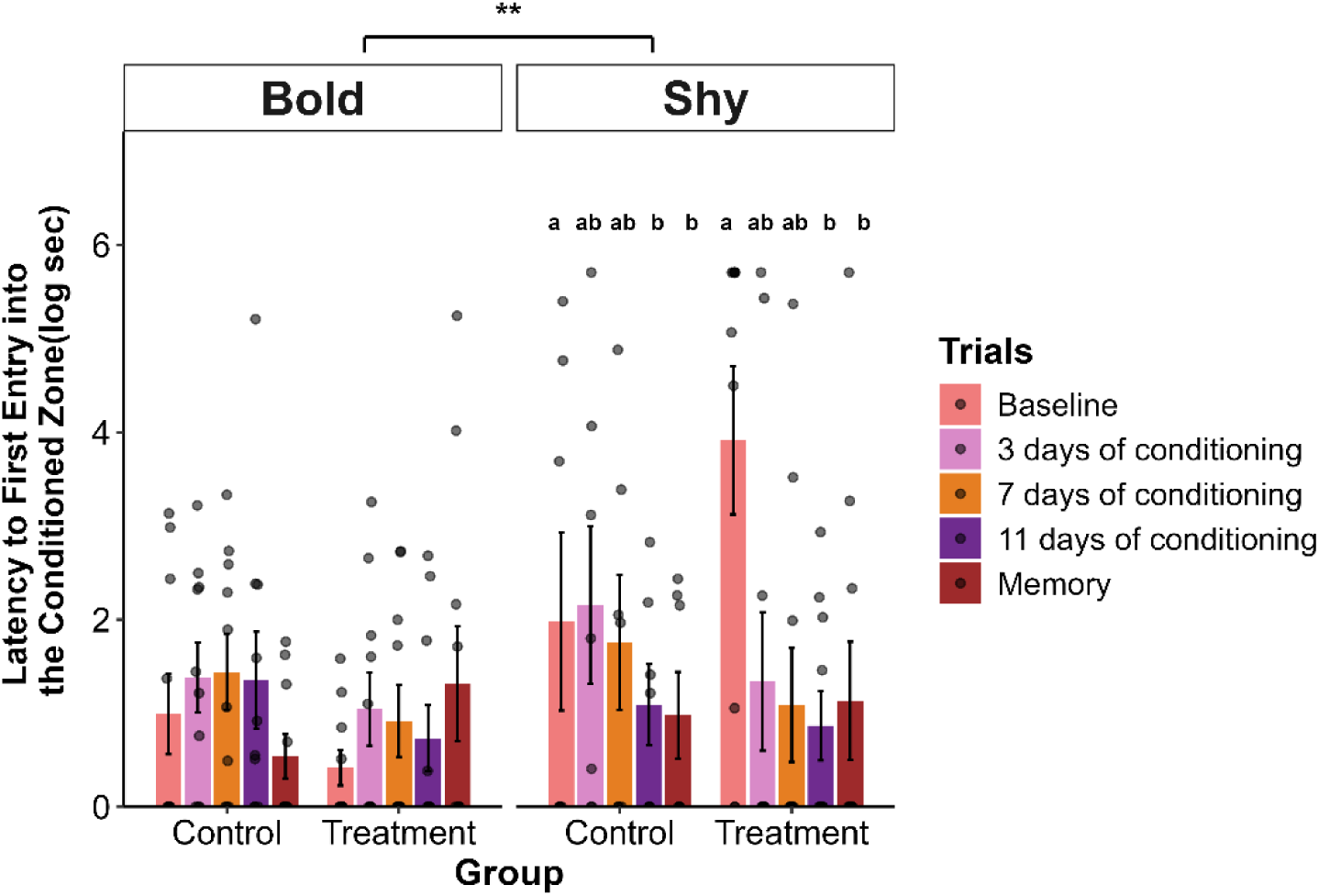
Latency to first entry into the conditioned zone across personality, treatment, and conditioned day. Peach bars are at baseline, pink bars represent learning after 3 days of conditioning sessions (Probe 1), orange bars are after 7 days of conditioning sessions (Probe 2), and purple bars are after 11 days of conditioning sessions (Probe 3). Red bars indicate memory retention seven days after the final conditioning session. Asterisk represents main effect of personality. Bars represent the mean latency to first entry time into the conditioned zone during the final 5 min of each trial for bold and shy zebrafish in the control and treatment groups at each trial for bold and shy zebrafish in the control and treatment treatments at each conditioned days. Error bars indicate SEM, and individual points represent individual fish. Different letters represent post hoc comparison within the shy group.

There was a significant personality by conditioned day interaction effect ( *F*_3.69,121.69_=2.941, p = 0.027, *n*^2^*G* = 0.067) on latency to first enter the CZ. Bold fish had significantly shorter latency to first entry only at baseline (p = 0.001) relative to the shy group. Only within the shy group, the latency to first entry significantly decreased at day 11 post-conditioning (p = 0.011) relative to baseline (Fig.3). Within the bold group, we did not observe any differences between baseline and probes in latency to first into the conditioned zone. See supplementary table 3 for full post-hoc comparisons.

For the memory trial, only within the shy group we observed a significant decrease in latency to first entry into the CZ when compared to baseline (p = 0.033) (Fig. 3), but this was not seen within the bold group between baseline and probes in latency to first into the conditioned zone (Supplementary Table 3). Overall, there was no significant difference between the bold and shy group for latency to first into the CZ (p = 0.811). See Supplementary Table 3 for full post-hoc comparisons.

## 4. Discussion

Understanding how animals retain learned associations is important for explaining how individuals adapt to changing environments. In this study, we examined how personality, food reward-based conditioning, sex, and number of conditioning days influence associative learning and memory retention in zebrafish using a conditioned place preference (CPP) paradigm. Overall, our findings suggest that shy fish have capabilities to associate and retain conditioning with food rewards for at least 7 days. Personalities differed in how they associate with the positive stimulus, with bold fish showing greater early engagement in the CZ and shy fish showing progressively stronger conditioned-zone responses across days of conditioning. These findings support the predictions that personality, sex, and treatment shape cognition across days of conditioning.

One framework linking personality and cognition involves a risk-reward trade off where bold personality have greater potential to find resources by taking higher risks and performing quickly. In our study, we found that the bold fish displayed stronger early conditioned zone engagement than the shy fish as indicated by more time spent in the conditioned zone at baseline and day 3 post conditioning, entered it more frequently, and exhibited shorter latency to first entry at baseline. This pattern suggests a reflection of their naturally exploratory, high-risk phenotype [6,23,42,52–55], which increases the likelihood of entering and sampling a novel or reward-associated zone, rather than a learning response [16]. We observed the shy fish showed progressive engagement into the conditioned zone by spending more time in the conditioned zone after 7 days of conditioning relative to baseline. At the same time, frequency and entry tendency became evident 11 days after conditioning. Our findings are consistent with studies in teleost fish, lizards, and birds that show personality can affect behavioral responses and flexibility during learning tasks [16,24,51–53]. For example, in a brook trout study, shy individuals were increasingly more successful in associating food cues with learning than bold individuals [16]. This may reflect differences in behavioral flexibility, as bold individuals have been reported to be less flexible when adapting to novel situations. Together, these findings suggest that personality may shape the response of reward-associated learning.

Reduced engagement with novel stimuli is commonly associated with shy behavioral phenotypes, and this trait in addition to genetic factors can limit the rate of learning [6,41]. Studies have reported food reward as a positive motivational stimulus that increases task engagement[14]. For example, a zebrafish study reported that food rewards decreased inhibitory avoidance learning by decreasing cnr1 expression levels [54]. Although we found no overall main effect of treatment, treatment effects depended on the number of conditioned days as indicated by the treatment by conditioned day interaction effects. Within the treatment group, time spent in the CZ and frequency of entry showed an increase after day 7 and 11 relative to baseline. Additionally, we found personality interactive effects where within the shy-treated fish, there was increased in time spent in the conditioned zone across conditioning compared to baseline. Similarly, within the shy treated, we found a decreased latency to first entry compared to shy control and bold groups. However, bold-treated fish overall entered the conditioned zone more frequently but showed little change in behavior from baseline relative to shy treated. Our findings contrast with a previous CPP study showing bold-treated males increased time spent in the CZ relative to baseline [6]. This discrepancy may reflect sex-dependent responses or cohort and natural variation within the lines. Together, our findings suggest that food rewards influenced interactions with the conditioned zone for both personalities, but may be more important for motivating the shy fish by reducing behavioral inhibition.

Sex differences have also been reported in cognition [24,33,55,56]. We found similar effects in sex-related patterns, which primarily influenced frequency of entry, where shy-treated males showed a progressive increase in conditioned zone entry at day 3, 7, and 11 post-conditioning. This suggests sex-dependent differences in exploratory strategy, anxiety-like behavior, and reward-associated task engagement. Previous studies have reported that female zebrafish can be bolder than males [30] and also show higher anxiety-like behavior depending on the task [34], indicating that sex effects on cognition are context-dependent. Additionally, other studies in fish have reported that males and females can differ in spatial exploration and learning strategies, with these differences shaped by ecological context, habitat, and task demands [30,56]. Because sex differences have been reported inconsistently in reward-associated cognition [6,34,57], this pattern may reflect context-dependent differences in motivation, exploration, and stress response potentially shaped by hormonal and genetic factors [58,59].

A major goal of this study was to examine personality variations in memory retention. We found no significant personality difference seven days after the last conditioning. However, because bold fish showed early zone engagement, which remained consistent for all subsequent probe trials, it is unclear if learning or recall occurred. However, shy-treated fish had reduced latency to first entry and increased time spent in the conditioned zone during the memory probe relative to baseline. Performance during the memory probe was similar to performance after 11 days of conditioning. These patterns suggest that behavioral changes acquired across repeated food-reward conditioning persisted after conditioning ceased. Previous appetitive studies have shown that shy or less exploratory individuals can retain food-associated learning across different intervals such as 24 hours and 3 days [30,60]. Our findings extend this by showing that conditioned-related behavioral changes in the shy fish remain for at least 7 days. These results suggest that personality-specific memory may depend on the type of task and stimulus valence. We also found sex specific patterns where only shy-treated males had significantly higher conditioned zone entries in the memory trial compared to baseline and after 3 days of conditioning. This sex-specific pattern is noteworthy given a previous study on zebrafish reported that males display higher memory retention than females [30]. However, in our study, it may not indicate a general male advantage but may reflect context-dependent differences in the nature of the task and cognitive strategy between the sexes [55].

In conclusion, this study found evidence that personality, food reward, and sex differences influence changes in conditioned behavior associated with cognition. Our findings demonstrate that a shy personality and male sex may require repeated exposure to rewards to reduce neophobia to the conditioned zones. We also show that shy fish are able to recall learned associations up to 7 days after conditioning. Collectively, our findings suggest that personality and sex related differences in learning and memory may depend on the task type and valence of the conditioning stimulus, making them employ different cognitive strategies in response to reward-associated memory. Future studies can explore longer memory retention intervals and underlying neurobiological mechanisms to highlight how personality and sex differences in associative learning.

## Acknowledgements

We thank Beth Trail, Brandon Wolfsohn and Amanda Hosert for help with fish husbandry. We also thank Jamie Corcoran for feedback on an earlier version of the manuscript.

## Author contributions

PSJ: Cognition assay development, data acquisition, analyses, and interpretation, manuscript drafting and editing. TTAL: Data acquisition, data analysis, manuscript editing. RYW: Project conceptualization and oversight; funding acquisition; data interpretation; manuscript editing.

## Declarations

### Animal ethics and consent to participate

Ethical approval for this study was obtained from the University of Nebraska at Omaha’s Institutional Animal Care and Use Committee, IACUC. The present study followed national and institutional guidelines for humane animal treatment and complied with relevant legislation from IACUC.

