## Supplemental Tables for "Personality Variation in Associative Learning and Memory Retention in Zebrafish (*Danio rerio*)"

Tran Thien An Le[1]

*Ryan Y. Wong[1],

[1]University of Nebraska at Omaha Biology Department, Omaha, NE USA

#### 1. Supplementary Tables: Time Spent in the Conditioned Zone

**Repeated measures ANOVA for time spent in the conditioned zone.**

| Effect | DFn | DFd | F | p | p<.05 | ges |
|---|---|---|---|---|---|---|
| Treatment | 1.00 | 33.00 | 2.361 | 0.134 | | 0.014 |
| Personality | 1.00 | 33.00 | 8.522 | 0.006 | * | 0.050 |
| Conditioned days | 2.91 | 96.14 | 4.314 | 0.007 | * | 0.094 |
| Treatment: personality | 1.00 | 33.00 | 2.449 | 0.127 | | 0.015 |
| Treatment: Conditioned days | 2.91 | 96.14 | 3.767 | 0.014 | * | 0.083 |
| Personality: Conditioned days | 2.91 | 96.14 | 3.220 | 0.027 | * | 0.072 |
| Treatment: Personality: Conditioned days | 2.91 | 96.14 | 2.259 | 0.088 | | 0.052 |

**Post hoc comparisons for treatment within each conditioned day.**

| contrast | conditioned days | estimate | SE | df | t.ratio | p.value |
|---|---|---|---|---|---|---|
| Control - Treatment | Bas | 1.28090771 | 0.4150715 | 33 | 3.08599270 | 0.004088945 |
| Control - Treatment | Pro.1 | 0.04179106 | 0.4202909 | 33 | 0.09943366 | 0.921395558 |
| Control - Treatment | Pro.2 | -0.33991500 | 0.2665324 | 33 | -1.27532346 | 0.211100154 |
| Control - Treatment | Pro.3 | -0.14897043 | 0.1509369 | 33 | -0.98697185 | 0.330838122 |
| Control - Treatment | Pro.4 | 0.30689523 | 0.3098392 | 33 | 0.99049829 | 0.329138124 |

**Post hoc comparisons for conditioned day within each treatment group.**

| contrast | treatment | estimate | SE | df | t.ratio | p.value |
|---|---|---|---|---|---|---|
| Bas - Pro.1 | Control | 0.21520907 | 0.3974887 | 33 | 0.5414218 | 0.98224266168 |
| Bas - Pro.2 | Control | 0.16869101 | 0.3617292 | 33 | 0.4663462 | 0.98984300362 |
| Bas - Pro.3 | Control | -0.15047117 | 0.3225670 | 33 | -0.4664804 | 0.98983195620 |
| Bas - Pro.4 | Control | -0.24426096 | 0.4376453 | 33 | -0.5581254 | 0.98013307588 |
| Pro.1 - Pro.2 | Control | -0.04651806 | 0.3659724 | 33 | -0.1271081 | 0.99993780376 |
| Pro.1 - Pro.3 | Control | -0.36568024 | 0.3306277 | 33 | -1.1060180 | 0.80216013595 |
| Pro.1 - Pro.4 | Control | -0.45947003 | 0.4114281 | 33 | -1.1167688 | 0.79657811111 |
| Pro.2 - Pro.3 | Control | -0.31916218 | 0.2265582 | 33 | -1.4087424 | 0.62655104780 |
| Pro.2 - Pro.4 | Control | -0.41295197 | 0.2627372 | 33 | -1.5717300 | 0.52525383396 |
| Pro.3 - Pro.4 | Control | -0.09378979 | 0.2402597 | 33 | -0.3903685 | 0.99484401881 |
| Bas - Pro.1 | Treatment | -1.02390759 | 0.3607151 | 33 | -2.8385490 | 0.05549572009 |
| Bas - Pro.2 | Treatment | -1.45213170 | 0.3282639 | 33 | -4.4236719 | 0.00089030387 |

| contrast | treatment | estimate | SE | df | t.ratio | p.value |
|---|---|---|---|---|---|---|
| Bas - Pro.3 | Treatment | -1.58034931 | 0.2927248 | 33 | -5.3987548 | 0.00005318255 |
| Bas - Pro.4 | Treatment | -1.21827344 | 0.3971566 | 33 | -3.0674887 | 0.03254813803 |
| Pro.1 - Pro.2 | Treatment | -0.42822411 | 0.3321146 | 33 | -1.2893867 | 0.69938136533 |
| Pro.1 - Pro.3 | Treatment | -0.55644172 | 0.3000398 | 33 | -1.8545598 | 0.36101322457 |
| Pro.1 - Pro.4 | Treatment | -0.19436585 | 0.3733649 | 33 | -0.5205788 | 0.98465203264 |
| Pro.2 - Pro.3 | Treatment | -0.12821761 | 0.2055982 | 33 | -0.6236318 | 0.97021369309 |
| Pro.2 - Pro.4 | Treatment | 0.23385826 | 0.2384301 | 33 | 0.9808251 | 0.86193863154 |
| Pro.3 - Pro.4 | Treatment | 0.36207587 | 0.2180321 | 33 | 1.6606541 | 0.47116152500 |

**Post hoc comparisons for the main effect of conditioned day.**

| contrast | estimate | SE | df | t.ratio | p.value |
|---|---|---|---|---|---|
| Bas - Pro.1 | -0.40434926 | 0.2683807 | 33 | -1.5066260 | 0.565612678 |
| Bas - Pro.2 | -0.64172035 | 0.2442361 | 33 | -2.6274586 | 0.088266261 |
| Bas - Pro.3 | -0.86541024 | 0.2177942 | 33 | -3.9735228 | 0.003126342 |
| Bas - Pro.4 | -0.73126720 | 0.2954940 | 33 | -2.4747279 | 0.121180149 |
| Pro.1 - Pro.2 | -0.23737109 | 0.2471012 | 33 | -0.9606231 | 0.870602872 |
| Pro.1 - Pro.3 | -0.46106098 | 0.2232367 | 33 | -2.0653454 | 0.258910148 |
| Pro.1 - Pro.4 | -0.32691794 | 0.2777924 | 33 | -1.1768428 | 0.764252967 |
| Pro.2 - Pro.3 | -0.22368989 | 0.1529700 | 33 | -1.4623123 | 0.593227041 |
| Pro.2 - Pro.4 | -0.08954685 | 0.1773977 | 33 | -0.5047802 | 0.986320000 |
| Pro.3 - Pro.4 | 0.13414304 | 0.1622211 | 33 | 0.8269151 | 0.920362025 |

**Post hoc comparisons for personality across each conditioned day.**

| contrast | conditioned days | estimate | SE | df | t.ratio | p.value |
|---|---|---|---|---|---|---|
| Bold - Shy | Bas | 1.3501425 | 0.4150715 | 33 | 3.2527947 | 0.002637402 |
| Bold - Shy | Pro.1 | 0.5734310 | 0.4202909 | 33 | 1.3643671 | 0.181684868 |
| Bold - Shy | Pro.2 | 0.2515893 | 0.2665324 | 33 | 0.9439350 | 0.352062966 |
| Bold - Shy | Pro.3 | -0.2320654 | 0.1509369 | 33 | -1.5375002 | 0.133705059 |
| Bold - Shy | PRO.4 | 0.2239329 | 0.3098392 | 33 | 0.7227391 | 0.474931564 |

**Post hoc comparisons for conditioned day within each personality type.**

| contrast | personality | estimate | SE | df | t.ratio | p.value |
|---|---|---|---|---|---|---|
| Bas - Pro.1 | Bold | -0.01599354 | 0.3607151 | 33 | -0.04433841 | 0.9999990723 |
| Bas - Pro.2 | Bold | -0.09244373 | 0.3282639 | 33 | -0.28161408 | 0.9985498353 |
| Bas - Pro.3 | Bold | -0.07430627 | 0.2927248 | 33 | -0.25384347 | 0.9990352495 |
| Bas - Pro.4 | Bold | -0.16816242 | 0.3971566 | 33 | -0.42341589 | 0.9929614207 |
| Pro.1 - Pro.2 | Bold | -0.07645020 | 0.3321146 | 33 | -0.23019223 | 0.9993437072 |
| Pro.1 - Pro.3 | Bold | -0.05831274 | 0.3000398 | 33 | -0.19435002 | 0.9996638654 |
| Pro.1 - Pro.4 | Bold | -0.15216889 | 0.3733649 | 33 | -0.40756078 | 0.9939169535 |
| Pro.2 - Pro.3 | Bold | 0.01813746 | 0.2055982 | 33 | 0.08821796 | 0.9999855056 |
| Pro.2 - Pro.4 | Bold | -0.07571869 | 0.2384301 | 33 | -0.31757181 | 0.9976808859 |
| Pro.3 - Pro.4 | Bold | -0.09385615 | 0.2180321 | 33 | -0.43046945 | 0.9925037261 |
| Bas - Pro.1 | Shy | -0.79270499 | 0.3974887 | 33 | -1.99428294 | 0.2910032921 |
| Bas - Pro.2 | Shy | -1.19099696 | 0.3617292 | 33 | -3.29251010 | 0.0187583017 |
| Bas - Pro.3 | Shy | -1.65651421 | 0.3225670 | 33 | -5.13541128 | 0.0001146713 |
| Bas - Pro.4 | Shy | -1.29437198 | 0.4376453 | 33 | -2.95758239 | 0.0422051829 |
| Pro.1 - Pro.2 | Shy | -0.39829198 | 0.3659724 | 33 | -1.08831140 | 0.8112097198 |
| Pro.1 - Pro.3 | Shy | -0.86380922 | 0.3306277 | 33 | -2.61263391 | 0.0910895828 |
| Pro.1 - Pro.4 | Shy | -0.50166699 | 0.4114281 | 33 | -1.21933097 | 0.7403462586 |
| Pro.2 - Pro.3 | Shy | -0.46551724 | 0.2265582 | 33 | -2.05473553 | 0.2635459045 |
| Pro.2 - Pro.4 | Shy | -0.10337502 | 0.2627372 | 33 | -0.39345402 | 0.9946856712 |
| Pro.3 - Pro.4 | Shy | 0.36214223 | 0.2402597 | 33 | 1.50729523 | 0.5651960396 |

**Post hoc comparisons for conditioned day within each treatment group × personality combination.**

| contrast | treatment | personality | estimate | SE | df | t.ratio | p.value |
|---|---|---|---|---|---|---|---|
| Bas - Pro 1 | Control | Bold | 0.064095 | 0.439926 | 132 | 0.1456965 | 0.99989741496 |
| Bas - Pro 2 | Control | Bold | 0.102447 | 0.439926 | 132 | 0.23287376 | 0.99934097767 |
| Bas - Pro 3 | Control | Bold | 0.113127 | 0.439926 | 132 | 0.25715044 | 0.99902569386 |
| Bas - Pro 4 | Control | Bold | -0.14559 | 0.439926 | 132 | -0.3309619 | 0.99738145008 |
| Pro 1 - Pro 2 | Control | Bold | 0.038351 | 0.439926 | 132 | 0.08717726 | 0.99998676443 |
| Pro 1 - Pro 3 | Control | Bold | 0.049031 | 0.439926 | 132 | 0.11145394 | 0.99996472188 |

| contrast | treatment | personality | estimate | SE | df | t.ratio | p.value |
|---|---|---|---|---|---|---|---|
| Pro 1 - Pro 4 | Control | Bold | -0.20969 | 0.439926 | 132 | -0.4766584 | 0.98934502133 |
| Pro 2 - Pro 3 | Control | Bold | 0.010679 | 0.439926 | 132 | 0.02427668 | 0.99999992013 |
| Pro 2 - Pro 4 | Control | Bold | -0.24804 | 0.439926 | 132 | -0.5638357 | 0.98000953539 |
| Pro 3 - Pro 4 | Control | Bold | -0.25872 | 0.439926 | 132 | -0.5881124 | 0.97664522875 |
| Bas - Pro 1 | Treatment | Bold | -0.09608 | 0.439926 | 132 | -0.2184065 | 0.99948850910 |
| Bas - Pro 2 | Treatment | Bold | -0.28733 | 0.439926 | 132 | -0.6531427 | 0.96578216302 |
| Bas - Pro 3 | Treatment | Bold | -0.26173 | 0.439926 | 132 | -0.5949626 | 0.97563030920 |
| Bas - Pro 4 | Treatment | Bold | -0.19072 | 0.439926 | 132 | -0.4335403 | 0.99257161018 |
| Pro 1 - Pro 2 | Treatment | Bold | -0.19125 | 0.439926 | 132 | -0.4347362 | 0.99249300206 |
| Pro 1 - Pro 3 | Treatment | Bold | -0.16565 | 0.439926 | 132 | -0.3765561 | 0.99567882359 |
| Pro 1 - Pro 4 | Treatment | Bold | -0.09464 | 0.439926 | 132 | -0.2151338 | 0.99951815715 |
| Pro 2 - Pro 3 | Treatment | Bold | 0.025594 | 0.439926 | 132 | 0.05818008 | 0.99999736908 |
| Pro 2 - Pro 4 | Treatment | Bold | 0.096608 | 0.439926 | 132 | 0.21960236 | 0.99947734579 |
| Pro 3 - Pro 4 | Treatment | Bold | 0.071013 | 0.439926 | 132 | 0.16142228 | 0.99984578206 |
| Bas - Pro 1 | Control | Shy | 0.366322 | 0.525812 | 132 | 0.69667840 | 0.95689347513 |
| Bas - Pro 2 | Control | Shy | 0.234934 | 0.525812 | 132 | 0.44680294 | 0.99166622108 |
| Bas - Pro 3 | Control | Shy | -0.41406 | 0.525812 | 132 | -0.7874849 | 0.93385007774 |
| Bas - Pro 4 | Control | Shy | -0.34292 | 0.525812 | 132 | -0.6521770 | 0.96596419975 |
| Pro 1 - Pro 2 | Control | Shy | -0.13138 | 0.525812 | 132 | -0.2498754 | 0.99912982456 |
| Pro 1 - Pro 3 | Control | Shy | -0.78039 | 0.525812 | 132 | -1.4841633 | 0.57456339050 |
| Pro 1 - Pro 4 | Control | Shy | -0.70924 | 0.525812 | 132 | -1.3488554 | 0.66123492870 |
| Pro 2 - Pro 3 | Control | Shy | -0.64900 | 0.525812 | 132 | -1.2342872 | 0.73137526746 |
| Pro 2 - Pro 4 | Control | Shy | -0.57785 | 0.525812 | 132 | -1.0989799 | 0.80678888679 |
| Pro 3 - Pro 4 | Control | Shy | 0.071146 | 0.525812 | 132 | 0.13530798 | 0.99992358305 |
| Bas - Pro 1 | Treatment | Shy | -1.95173 | 0.439926 | 132 | -4.4364964 | 0.00018348668 |
| Bas - Pro 2 | Treatment | Shy | -2.61692 | 0.439926 | 132 | -5.9485586 | 0.00000022737 |
| Bas - Pro 3 | Treatment | Shy | -2.89895 | 0.439926 | 132 | -6.5896434 | 0.00000000958 |
| Bas - Pro 4 | Treatment | Shy | -2.24582 | 0.439926 | 132 | -5.1049913 | 0.00001108999 |
| Pro 1 - Pro 2 | Treatment | Shy | -0.66519 | 0.439926 | 132 | -1.5120622 | 0.55653670768 |
| Pro 1 - Pro 3 | Treatment | Shy | -0.94722 | 0.439926 | 132 | -2.1531470 | 0.20427466051 |
| Pro 1 - Pro 4 | Treatment | Shy | -0.29408 | 0.439926 | 132 | -0.6684949 | 0.96280115857 |

| contrast | treatment | personality | estimate | SE | df | t.ratio | p.value |
|---|---|---|---|---|---|---|---|
| Pro 2 - Pro 3 | Treatment | Shy | -0.28203 | 0.439926 | 132 | -0.6410846 | 0.96800904152 |
| Pro 2 - Pro 4 | Treatment | Shy | 0.371107 | 0.439926 | 132 | 0.84356733 | 0.91644167335 |
| Pro 3 - Pro 4 | Treatment | Shy | 0.653137 | 0.439926 | 132 | 1.48465208 | 0.57424756854 |

### Personality Variation in Associative Learning and Memory Retention in Zebrafish (*Danio rerio*)

Princess Sunday-Jimmy[1]

Tran Thien An Le[1]

*Ryan Y. Wong[1],

[1]University of Nebraska at Omaha Biology Department, Omaha, NE USA

#### 2. Supplementary Tables: Frequency of Entry into the Conditioned Zone

**Repeated-measures ANOVA for frequency of entry into the conditioned zone.**

| Effect | DFn | DFd | F | p | p<.05 | ges |
|---|---|---|---|---|---|---|
| Treatment | 1.00 | 29.00 | 0.486 | 0.491 | | 0.005000 |
| Personality | 1.00 | 29.00 | 12.904 | 0.001 | * | 0.117000 |
| Sex | 1.00 | 29.00 | 8.844 | 0.006 | * | 0.083000 |
| Conditioned days | 3.33 | 96.59 | 1.249 | 0.296 | | 0.029000 |
| Treatment: personality | 1.00 | 29.00 | 3.490 | 0.072 | | 0.035000 |
| Treatment: Sex | 1.00 | 29.00 | 0.035 | 0.853 | | 0.000359 |
| Personality: Sex | 1.00 | 29.00 | 3.325 | 0.079 | | 0.033000 |
| Treatment: Conditioned days | 3.33 | 96.59 | 3.139 | 0.025 | * | 0.071000 |
| Personality: Conditioned days | 3.33 | 96.59 | 3.604 | 0.013 | * | 0.080000 |
| Sex: Conditioned days | 3.33 | 96.59 | 3.680 | 0.012 | * | 0.082000 |
| Treatment: personality: sex | 1.00 | 29.00 | 0.960 | 0.335 | | 0.010000 |
| Treatment: personality: Conditioned days | 3.33 | 96.59 | 1.641 | 0.180 | | 0.038000 |
| Treatment: Sex:Conditioned days | 3.33 | 96.59 | 1.723 | 0.162 | | 0.040000 |
| personality:Sex:Conditioned days | 3.33 | 96.59 | 1.223 | 0.306 | | 0.029000 |
| Treatment: Personality: Sex: Conditioned days | 3.33 | 96.59 | 2.227 | 0.083 | | 0.051000 |

**Post hoc comparisons for treatment across each conditioned day.**

| contrast | conditioned days | estimate | SE | df | t.ratio | p.value |
|---|---|---|---|---|---|---|
| Control - Treatment | Bas | 0.5732963 | 0.1781401 | 29 | 3.2182323 | 0.00316729 |
| Control - Treatment | Pro.1 | 0.2442268 | 0.2598226 | 29 | 0.9399751 | 0.35499445 |
| Control - Treatment | Pro.2 | -0.2172414 | 0.2072824 | 29 | -1.0480458 | 0.30327411 |
| Control - Treatment | Pro.3 | -0.3408047 | 0.2139193 | 29 | -1.5931464 | 0.12197005 |
| Control - Treatment | Pro.4 | 0.1625007 | 0.2410862 | 29 | 0.6740359 | 0.50562613 |

**Post hoc comparisons for conditioned day within each treatment group.**

| contrast | treatment | estimate | SE | df | t.ratio | p.value |
|---|---|---|---|---|---|---|
| Bas - Pro.1 | Control | 0.247213498 | 0.2367168 | 29 | 1.04434274 | 0.83263648 |
| Bas - Pro.2 | Control | 0.233704921 | 0.1913055 | 29 | 1.22163185 | 0.73907766 |
| Bas - Pro.3 | Control | 0.370227115 | 0.1903918 | 29 | 1.94455358 | 0.31763474 |
| Bas - Pro.4 | Control | 0.323519618 | 0.2215448 | 29 | 1.46028982 | 0.59528990 |
| Pro.1 - Pro.2 | Control | -0.013508576 | 0.2232366 | 29 | -0.06051236 | 0.99999676 |
| Pro.1 - Pro.3 | Control | 0.123013617 | 0.2490129 | 29 | 0.49400491 | 0.98730565 |
| Pro.1 - Pro.4 | Control | 0.076306121 | 0.2771100 | 29 | 0.27536401 | 0.99866226 |
| Pro.2 - Pro.3 | Control | 0.136522193 | 0.1567905 | 29 | 0.87072979 | 0.90526143 |

| contrast | treatment | estimate | SE | df | t.ratio | p.value |
|---|---|---|---|---|---|---|
| Pro.2 - Pro.4 | Control | 0.089814697 | 0.1893745 | 29 | 0.47427025 | 0.98910886 |
| Pro.3 - Pro.4 | Control | -0.046707496 | 0.2303609 | 29 | -0.20275788 | 0.99959950 |
| Bas - Pro.1 | Treatment | -0.081855993 | 0.2116695 | 29 | -0.38671600 | 0.99499347 |
| Bas - Pro.2 | Treatment | -0.556832742 | 0.1710633 | 29 | -3.25512776 | 0.02227590 |
| Bas - Pro.3 | Treatment | -0.543873883 | 0.1702462 | 29 | -3.19463086 | 0.02573523 |
| Bas - Pro.4 | Treatment | -0.087275910 | 0.1981029 | 29 | -0.44055853 | 0.99176014 |
| Pro.1 - Pro.2 | Treatment | -0.474976749 | 0.1996157 | 29 | -2.37945589 | 0.14974987 |
| Pro.1 - Pro.3 | Treatment | -0.462017890 | 0.2226646 | 29 | -2.07495011 | 0.25784408 |
| Pro.1 - Pro.4 | Treatment | -0.005419917 | 0.2477886 | 29 | -0.02187314 | 0.99999994 |
| Pro.2 - Pro.3 | Treatment | 0.012958859 | 0.1402003 | 29 | 0.09243101 | 0.99998240 |
| Pro.2 - Pro.4 | Treatment | 0.469556832 | 0.1693366 | 29 | 2.77292054 | 0.06713283 |
| Pro.3 - Pro.4 | Treatment | 0.456597973 | 0.2059862 | 29 | 2.21664387 | 0.20211886 |

**Post hoc comparisons for personality across each conditioned day.**

| contrast | timepoint | estimate | SE | df | t.ratio | p.value |
|---|---|---|---|---|---|---|
| Bold - Shy | Bas | 0.88236208 | 0.1781401 | 29 | 4.95319135 | 0.000028891 |
| Bold - Shy | Pro.1 | 0.81607219 | 0.2598226 | 29 | 3.14088215 | 0.003857504 |
| Bold - Shy | Pro.2 | 0.23787909 | 0.2072824 | 29 | 1.14760896 | 0.260509572 |
| Bold - Shy | Pro.3 | -0.02059136 | 0.2139193 | 29 | -0.09625762 | 0.923978100 |
| Bold - Shy | Pro.4 | 0.25890830 | 0.2410862 | 29 | 1.07392427 | 0.291711562 |

**Post hoc comparisons for conditioned day within each personality type.**

| contrast | personality | estimate | SE | df | t.ratio | p.value |
|---|---|---|---|---|---|---|
| Bas - Pro.1 | Bold | 0.11582370 | 0.2044922 | 29 | 0.5663967 | 0.97891106 |
| Bas - Pro.2 | Bold | 0.16067758 | 0.1652628 | 29 | 0.9722550 | 0.86537671 |
| Bas - Pro.3 | Bold | 0.36465333 | 0.1644735 | 29 | 2.2170949 | 0.20195687 |
| Bas - Pro.4 | Bold | 0.42984874 | 0.1913856 | 29 | 2.2459832 | 0.19178067 |
| Pro.1 - Pro.2 | Bold | 0.04485389 | 0.1928471 | 29 | 0.2325878 | 0.99931125 |
| Pro.1 - Pro.3 | Bold | 0.24882964 | 0.2151144 | 29 | 1.1567316 | 0.77519626 |
| Pro.1 - Pro.4 | Bold | 0.31402504 | 0.2393866 | 29 | 1.3117905 | 0.68623456 |
| Pro.2 - Pro.3 | Bold | 0.20397575 | 0.1354464 | 29 | 1.5059519 | 0.56701668 |

| contrast | personality | estimate | SE | df | t.ratio | p.value |
|---|---|---|---|---|---|---|
| Pro.2 - Pro.4 | Bold | 0.26917116 | 0.1635947 | 29 | 1.6453542 | 0.48192356 |
| Pro.3 - Pro.4 | Bold | 0.06519541 | 0.1990015 | 29 | 0.3276126 | 0.99736368 |
| Bas - Pro.1 | Shy | 0.04953381 | 0.2429440 | 29 | 0.2038898 | 0.99959058 |
| Bas - Pro.2 | Shy | -0.48380540 | 0.1963381 | 29 | -2.4641444 | 0.12711390 |
| Bas - Pro.3 | Shy | -0.53830010 | 0.1954004 | 29 | -2.7548572 | 0.06980116 |
| Bas - Pro.4 | Shy | -0.19360503 | 0.2273729 | 29 | -0.8514870 | 0.91195562 |
| Pro.1 - Pro.2 | Shy | -0.53333921 | 0.2291092 | 29 | -2.3278821 | 0.16503865 |
| Pro.1 - Pro.3 | Shy | -0.58783391 | 0.2555636 | 29 | -2.3001474 | 0.17375066 |
| Pro.1 - Pro.4 | Shy | -0.24313884 | 0.2843998 | 29 | -0.8549192 | 0.91078221 |
| Pro.2 - Pro.3 | Shy | -0.05449470 | 0.1609151 | 29 | -0.3386549 | 0.99700145 |
| Pro.2 - Pro.4 | Shy | 0.29020037 | 0.1943563 | 29 | 1.4931361 | 0.57494710 |
| Pro.3 - Pro.4 | Shy | 0.34469507 | 0.2364209 | 29 | 1.4579720 | 0.59672540 |

**Post hoc comparisons for sex across each conditioned day.**

| contrast | conditioned days | estimate | SE | df | t.ratio | p.value |
|---|---|---|---|---|---|---|
| F - M | Bas | 0.8962947 | 0.1781401 | 29 | 5.0314031 | 0.00002324588 |
| F - M | Pro.1 | 0.5700903 | 0.2598226 | 29 | 2.1941520 | 0.03638616688 |
| F - M | Pro.2 | 0.1908379 | 0.2072824 | 29 | 0.9206661 | 0.36482048441 |
| F - M | Pro.3 | 0.3050498 | 0.2139193 | 29 | 1.4260043 | 0.16454378293 |
| F - M | Pro.4 | -0.1620040 | 0.2410862 | 29 | -0.6719756 | 0.50691787395 |

**Post hoc comparisons for conditioned day within each sex.**

| contrast | sex | estimate | SE | df | t.ratio | p.value |
|---|---|---|---|---|---|---|
| Bas - Pro.1 | F | 0.24578097 | 0.2586644 | 29 | 0.9501924 | 0.87468304 |
| Bas - Pro.2 | F | 0.19116452 | 0.2090427 | 29 | 0.9144758 | 0.88900522 |
| Bas - Pro.3 | F | 0.20879906 | 0.2080443 | 29 | 1.0036277 | 0.85155720 |
| Bas - Pro.4 | F | 0.64727122 | 0.2420857 | 29 | 2.6737275 | 0.08295763 |
| Pro.1 - Pro.2 | F | -0.05461645 | 0.2439344 | 29 | -0.2238981 | 0.99940731 |
| Pro.1 - Pro.3 | F | -0.03698190 | 0.2721006 | 29 | -0.1359126 | 0.99991817 |
| Pro.1 - Pro.4 | F | 0.40149025 | 0.3028027 | 29 | 1.3259136 | 0.67774135 |
| Pro.2 - Pro.3 | F | 0.01763454 | 0.1713276 | 29 | 0.1029288 | 0.99997297 |

| contrast | sex | estimate | SE | df | t.ratio | p.value |
|---|---|---|---|---|---|---|
| Pro.2 - Pro.4 | F | 0.45610670 | 0.2069327 | 29 | 2.2041306 | 0.20665202 |
| Pro.3 - Pro.4 | F | 0.43847216 | 0.2517193 | 29 | 1.7419095 | 0.42542463 |
| Bas - Pro.1 | M | -0.08042346 | 0.1842052 | 29 | -0.4365972 | 0.99203818 |
| Bas - Pro.2 | M | -0.51429234 | 0.1488676 | 29 | -3.4546957 | 0.01369668 |
| Bas - Pro.3 | M | -0.38244583 | 0.1481566 | 29 | -2.5813618 | 0.10048108 |
| Bas - Pro.4 | M | -0.41102751 | 0.1723988 | 29 | -2.3841665 | 0.14841137 |
| Pro.1 - Pro.2 | M | -0.43386888 | 0.1737154 | 29 | -2.4975850 | 0.11898119 |
| Pro.1 - Pro.3 | M | -0.30202237 | 0.1937736 | 29 | -1.5586351 | 0.53453866 |
| Pro.1 - Pro.4 | M | -0.33060405 | 0.2156378 | 29 | -1.5331450 | 0.55022160 |
| Pro.2 - Pro.3 | M | 0.13184651 | 0.1220092 | 29 | 1.0806276 | 0.81488663 |
| Pro.2 - Pro.4 | M | 0.10326483 | 0.1473650 | 29 | 0.7007420 | 0.95466771 |
| Pro.3 - Pro.4 | M | -0.02858168 | 0.1792593 | 29 | -0.1594433 | 0.99984557 |

**Post hoc comparisons for conditioned day within each treatment group × personality × sex combination.**

| contrast | treatment | personality | sex | estimate | SE | df | t.ratio | p.value |
|---|---|---|---|---|---|---|---|---|
| Bas - Pro.1 | Control | Bold | F | 0.143866 | 0.448019 | 29 | 0.32111638484 | 0.99756 |
| Bas - Pro.2 | Control | Bold | F | 0.290244 | 0.362072 | 29 | 0.80161970439 | 0.92799 |
| Bas - Pro.3 | Control | Bold | F | 0.609784 | 0.360343 | 29 | 1.69223196024 | 0.45416 |
| Bas - Pro.4 | Control | Bold | F | 0.830012 | 0.419304 | 29 | 1.97949768322 | 0.30082 |
| Pro.1 - Pro.2 | Control | Bold | F | 0.146378 | 0.422506 | 29 | 0.34645131017 | 0.99672 |
| Pro.1 - Pro.3 | Control | Bold | F | 0.465918 | 0.471292 | 29 | 0.98859716569 | 0.85826 |
| Pro.1 - Pro.4 | Control | Bold | F | 0.686146 | 0.524469 | 29 | 1.30826664614 | 0.68834 |
| Pro.2 - Pro.3 | Control | Bold | F | 0.319539 | 0.296748 | 29 | 1.07680516912 | 0.81679 |
| Pro.2 - Pro.4 | Control | Bold | F | 0.539768 | 0.358417 | 29 | 1.50597426858 | 0.56700 |
| Pro.3 - Pro.4 | Control | Bold | F | 0.220228 | 0.435990 | 29 | 0.50512156810 | 0.9862 |
| Bas - Pro.1 | Treatment | Bold | F | 0.134953 | 0.448019 | 29 | 0.30122161215 | 0.9980 |
| Bas - Pro.2 | Treatment | Bold | F | 0.286600 | 0.362072 | 29 | 0.79155586785 | 0.9309 |
| Bas - Pro.3 | Treatment | Bold | F | 0.211051 | 0.360343 | 29 | 0.58569484594 | 0.9761 |
| Bas - Pro.4 | Treatment | Bold | F | 0.509861 | 0.419304 | 29 | 1.21596770001 | 0.7423 |
| Pro.1 - Pro.2 | Treatment | Bold | F | 0.151647 | 0.422506 | 29 | 0.35892309750 | 0.9962 |
| Pro.1 - Pro.3 | Treatment | Bold | F | 0.076097 | 0.471292 | 29 | 0.16146666714 | 0.9998 |

| contrast | treatment | personality | sex | estimate | SE | df | t.ratio | p.value |
|---|---|---|---|---|---|---|---|---|
| Pro.1 - Pro.4 | Treatment | Bold | F | 0.374907 | 0.524469 | 29 | 0.71483201225 | 0.9513 |
| Pro.2 - Pro.3 | Treatment | Bold | F | -0.07554 | 0.296748 | 29 | -0.25459123473 | 0.9990 |
| Pro.2 - Pro.4 | Treatment | Bold | F | 0.223260 | 0.358417 | 29 | 0.62290489982 | 0.9701 |
| Pro.3 - Pro.4 | Treatment | Bold | F | 0.298809 | 0.435990 | 29 | 0.685358411363 | 0.9580 |
| Bas - Pro.1 | Control | Shy | F | 0.312077 | 0.633595 | 29 | 0.492549214845 | 0.9874 |
| Bas - Pro.2 | Control | Shy | F | 0.245311 | 0.512048 | 29 | 0.479078966931 | 0.9886 |
| Bas - Pro.3 | Control | Shy | F | 0.071158 | 0.509602 | 29 | 0.139634548224 | 0.9999 |
| Bas - Pro.4 | Control | Shy | F | 0.312077 | 0.592986 | 29 | 0.526280401709 | 0.9839 |
| Pro.1 - Pro.2 | Control | Shy | F | -0.06676 | 0.597514 | 29 | -0.111738975681 | 0.9999 |
| Pro.1 - Pro.3 | Control | Shy | F | -0.24091 | 0.666507 | 29 | -0.361464784205 | 0.9961 |
| Pro.1 - Pro.4 | Control | Shy | F | 0.000000 | 0.741712 | 29 | 0.00000000 | 1.0000 |
| Pro.2 - Pro.3 | Control | Shy | F | -0.17415 | 0.419665 | 29 | -0.414981519374 | 0.9934 |
| Pro.2 - Pro.4 | Control | Shy | F | 0.066765 | 0.506879 | 29 | 0.131719064984 | 0.9999 |
| Pro.3 - Pro.4 | Control | Shy | F | 0.240919 | 0.616583 | 29 | 0.390732080837 | 0.9947 |
| Bas - Pro.1 | Treatment | Shy | F | 0.392226 | 0.517328 | 29 | 0.758177068129 | 0.9404 |
| Bas - Pro.2 | Treatment | Shy | F | -0.05749 | 0.418085 | 29 | -0.137528518529 | 0.9999 |
| Bas - Pro.3 | Treatment | Shy | F | -0.05679 | 0.416088 | 29 | -0.13650374340 | 0.9999 |
| Bas - Pro.4 | Treatment | Shy | F | 0.937133 | 0.484171 | 29 | 1.935541601508 | 0.3220 |
| Pro.1 - Pro.2 | Treatment | Shy | F | -0.44972 | 0.487868 | 29 | -0.921816568498 | 0.8861 |
| Pro.1 - Pro.3 | Treatment | Shy | F | -0.44902 | 0.544201 | 29 | -0.825107581133 | 0.9206 |
| Pro.1 - Pro.4 | Treatment | Shy | F | 0.544907 | 0.605605 | 29 | 0.899772331185 | 0.89462 |
| Pro.2 - Pro.3 | Treatment | Shy | F | 0.000701 | 0.342655 | 29 | 0.002045836601 | 0.99999 |
| Pro.2 - Pro.4 | Treatment | Shy | F | 0.994632 | 0.413865 | 29 | 2.403275603813 | 0.14307 |
| Pro.3 - Pro.4 | Treatment | Shy | F | 0.993931 | 0.503438 | 29 | 1.974285986384 | 0.30329 |
| Bas - Pro.1 | Control | Bold | M | 0.230084 | 0.365806 | 29 | 0.628978941842 | 0.96912 |
| Bas - Pro.2 | Control | Bold | M | 0.146330 | 0.295631 | 29 | 0.494975935368 | 0.98721 |
| Bas - Pro.3 | Control | Bold | M | 0.523345 | 0.294219 | 29 | 1.778762330741 | 0.40462 |
| Bas - Pro.4 | Control | Bold | M | 0.159332 | 0.342360 | 29 | 0.465392097798 | 0.98985 |
| Pro.1 - Pro.2 | Control | Bold | M | -0.08375 | 0.344975 | 29 | -0.242783923793 | 0.99918 |
| Pro.1 - Pro.3 | Control | Bold | M | 0.293261 | 0.384808 | 29 | 0.762096602616 | 0.93935 |
| Pro.1 - Pro.4 | Control | Bold | M | -0.07075 | 0.428227 | 29 | -0.16522211152 | 0.99982 |

| contrast | treatment | personality | sex | estimate | SE | df | t.ratio | p.value |
|---|---|---|---|---|---|---|---|---|
| Pro.2 - Pro.3 | Control | Bold | M | 0.377015 | 0.242293 | 29 | 1.556026195233 | 0.53614 |
| Pro.2 - Pro.4 | Control | Bold | M | 0.013001 | 0.292647 | 29 | 0.044428199381 | 0.99999 |
| Pro.3 - Pro.4 | Control | Bold | M | -0.36401 | 0.355984 | 29 | -1.02255450224 | 0.84289 |
| Bas - Pro.1 | Treatment | Bold | M | -0.04560 | 0.365806 | 29 | -0.124682756101 | 0.99994 |
| Bas - Pro.2 | Treatment | Bold | M | -0.08046 | 0.295631 | 29 | -0.272181331410 | 0.99872 |
| Bas - Pro.3 | Treatment | Bold | M | 0.114431 | 0.294219 | 29 | 0.388933350331 | 0.99488 |
| Bas - Pro.4 | Treatment | Bold | M | 0.220189 | 0.342360 | 29 | 0.643149088522 | 0.96655 |
| Pro.1 - Pro.2 | Treatment | Bold | M | -0.03485 | 0.344975 | 29 | -0.101037562811 | 0.99997 |
| Pro.1 - Pro.3 | Treatment | Bold | M | 0.160041 | 0.384808 | 29 | 0.415899020153 | 0.99338 |
| Pro.1 - Pro.4 | Treatment | Bold | M | 0.265798 | 0.428227 | 29 | 0.620695188986 | 0.97056 |
| Pro.2 - Pro.3 | Treatment | Bold | M | 0.194896 | 0.242293 | 29 | 0.804382242096 | 0.92715 |
| Pro.2 - Pro.4 | Treatment | Bold | M | 0.300654 | 0.292647 | 29 | 1.027361746734 | 0.84065 |
| Pro.3 - Pro.4 | Treatment | Bold | M | 0.105757 | 0.355984 | 29 | 0.297084237093 | 0.99819 |
| Bas - Pro.1 | Control | Shy | M | 0.302825 | 0.400721 | 29 | 0.755701297077 | 0.94108 |
| Bas - Pro.2 | Control | Shy | M | 0.252933 | 0.323847 | 29 | 0.781025858651 | 0.93406 |
| Bas - Pro.3 | Control | Shy | M | 0.276619 | 0.322300 | 29 | 0.858265959900 | 0.90962 |
| Bas - Pro.4 | Control | Shy | M | -0.00734 | 0.375037 | 29 | -0.019580749187 | 0.99999 |
| Pro.1 - Pro.2 | Control | Shy | M | -0.04989 | 0.377901 | 29 | -0.132024252634 | 0.99992 |
| Pro.1 - Pro.3 | Control | Shy | M | -0.02620 | 0.421536 | 29 | -0.062166990551 | 0.99999 |
| Pro.1 - Pro.4 | Control | Shy | M | -0.31016 | 0.469100 | 29 | -0.661200340971 | 0.96308 |
| Pro.2 - Pro.3 | Control | Shy | M | 0.023686 | 0.265419 | 29 | 0.089241771792 | 0.99998 |
| Pro.2 - Pro.4 | Control | Shy | M | -0.26027 | 0.320578 | 29 | -0.811896878336 | 0.92484 |
| Pro.3 - Pro.4 | Control | Shy | M | -0.28396 | 0.389961 | 29 | -0.728182651842 | 0.94816 |
| Bas - Pro.1 | Treatment | Shy | M | -0.80899 | 0.338671 | 29 | -2.388730565061 | 0.14712 |
| Bas - Pro.2 | Treatment | Shy | M | -2.37596 | 0.273701 | 29 | -8.680881671201 | 0.00000 |
| Bas - Pro.3 | Treatment | Shy | M | -2.44418 | 0.272394 | 29 | -8.972961716952 | 0.00000 |
| Bas - Pro.4 | Treatment | Shy | M | -2.01628 | 0.316964 | 29 | -6.361239426799 | 0.00000 |
| Pro.1 - Pro.2 | Treatment | Shy | M | -1.56697 | 0.319381 | 29 | -4.906219248892 | 0.00029 |
| Pro.1 - Pro.3 | Treatment | Shy | M | -1.63518 | 0.356263 | 29 | -4.589825241567 | 0.00070 |
| Pro.1 - Pro.4 | Treatment | Shy | M | -1.20729 | 0.396461 | 29 | -3.045169096689 | 0.03651 |
| Pro.2 - Pro.3 | Treatment | Shy | M | -0.06821 | 0.224320 | 29 | -0.304087119584 | 0.99802 |

| contrast | treatment | personality | sex | estimate | SE | df | t.ratio | p.value |
|---|---|---|---|---|---|---|---|---|
| Pro.2 - Pro.4 | Treatment | Shy | M | 0.359684 | 0.270938 | 29 | 1.327533991755 | 0.67676 |
| Pro.3 - Pro.4 | Treatment | Shy | M | 0.427893 | 0.329577 | 29 | 1.298306407265 | 0.69429 |

Personality Variation in Associative Learning and Memory Retention in Zebrafish (*Danio rerio*)

Princess Sunday-Jimmy[1]

Tran Thien An Le[1]

*Ryan Y. Wong[1],

[1]University of Nebraska at Omaha Biology Department, Omaha, NE USA

#### 3. Supplementary Tables: Latency to First Entry into the Conditioned Zone

**Repeated-measures ANOVA for latency to first entry into the conditioned zone.**

| Effect | DFn | DFd | F | p | p<.05 | ges |
|---|---|---|---|---|---|---|
| Treatment | 1.00 | 33.00 | 0.137 | 0.714 | | 0.000827 |
| Personality | 1.00 | 33.00 | 6.411 | 0.016 | * | 0.037000 |
| Conditioned day | 3.69 | 121.69 | 1.640 | 0.173 | | 0.038000 |
| Treatment: Personality | 1.00 | 33.00 | 0.465 | 0.500 | | 0.003000 |
| Treatment: Conditioned day | 3.69 | 121.69 | 1.265 | 0.289 | | 0.030000 |
| Personality: Conditioned day | 3.69 | 121.69 | 2.941 | 0.027 | * | 0.067000 |
| Treatment: Personality: Conditioned day | 3.69 | 121.69 | 1.380 | 0.247 | | 0.032000 |

**Post hoc comparisons for personality across each conditioned day.**

| contrast | conditioned day | estimate | SE | df | t.ratio | p.value |
|---|---|---|---|---|---|---|
| Bold - Shy | Bas | -2.24302243 | 0.6199260 | 33 | -3.6182098 | 0.0009800836 |
| Bold - Shy | Pro.1 | -0.53581266 | 0.5944974 | 33 | -0.9012868 | 0.3739670030 |
| Bold - Shy | Pro.2 | -0.24310303 | 0.5287165 | 33 | -0.4597984 | 0.6486759906 |
| Bold - Shy | Pro.3 | 0.06373997 | 0.4315554 | 33 | 0.1476983 | 0.8834796670 |
| Bold - Shy | Pro.4 | -0.12795585 | 0.5304761 | 33 | -0.2412095 | 0.8108852539 |

**Post hoc comparisons for conditioned day within each personality type.**

| contrast | personality | estimate | SE | df | t.ratio | p.value |
|---|---|---|---|---|---|---|
| Bas - Pro.1 | Bold | -0.50769370 | 0.5376444 | 33 | -0.94429275 | 0.87739319 |
| Bas - Pro.2 | Bold | -0.47386036 | 0.5577808 | 33 | -0.84954588 | 0.91289029 |
| Bas - Pro.3 | Bold | -0.33732967 | 0.5104900 | 33 | -0.66079579 | 0.96333718 |
| Bas - Pro.4 | Bold | -0.22189163 | 0.5610483 | 33 | -0.39549473 | 0.99457904 |
| Pro.1 - Pro.2 | Bold | 0.03383334 | 0.5165803 | 33 | 0.06549483 | 0.99999559 |
| Pro.1 - Pro.3 | Bold | 0.17036402 | 0.4976655 | 33 | 0.34232636 | 0.99689508 |
| Pro.1 - Pro.4 | Bold | 0.28580207 | 0.5672519 | 33 | 0.50383621 | 0.98641548 |
| Pro.2 - Pro.3 | Bold | 0.13653069 | 0.4959337 | 33 | 0.27530026 | 0.99867316 |
| Pro.2 - Pro.4 | Bold | 0.25196873 | 0.4216889 | 33 | 0.59752285 | 0.97449273 |
| Pro.3 - Pro.4 | Bold | 0.11543804 | 0.4964989 | 33 | 0.23250414 | 0.99931732 |
| Bas - Pro.1 | Shy | 1.19951607 | 0.5924553 | 33 | 2.02465257 | 0.27698843 |
| Bas - Pro.2 | Shy | 1.52605904 | 0.6146445 | 33 | 2.48283211 | 0.11921007 |

| contrast | personality | estimate | SE | df | t.ratio | p.value |
|---|---|---|---|---|---|---|
| Bas - Pro.3 | Shy | 1.96943273 | 0.5625326 | 33 | 3.50101065 | 0.01103155 |
| Bas - Pro.4 | Shy | 1.89317495 | 0.6182451 | 33 | 3.06217560 | 0.03296442 |
| Pro.1 - Pro.2 | Shy | 0.32654297 | 0.5692437 | 33 | 0.57364352 | 0.97802522 |
| Pro.1 - Pro.3 | Shy | 0.76991666 | 0.5484007 | 33 | 1.40393089 | 0.62953347 |
| Pro.1 - Pro.4 | Shy | 0.69365889 | 0.6250812 | 33 | 1.10971005 | 0.80025050 |
| Pro.2 - Pro.3 | Shy | 0.44337369 | 0.5464924 | 33 | 0.81130810 | 0.92528606 |
| Pro.2 - Pro.4 | Shy | 0.36711591 | 0.4646785 | 33 | 0.79004283 | 0.93169490 |
| Pro.3 - Pro.4 | Shy | -0.07625778 | 0.5471151 | 33 | -0.13938159 | 0.99991023 |

**Post hoc comparisons for conditioned day within each treatment × personality combination**.

| contrast | treatment | personality | estimate | SE | df | t.ratio | p.value |
|---|---|---|---|---|---|---|---|
| Bas - Pro.1 | Control | Bold | -0.38753379 | 0.7603440 | 33 | -0.50968219 | 1.000000000 |
| Bas - Pro.2 | Control | Bold | -0.44698912 | 0.7888211 | 33 | -0.56665459 | 1.000000000 |
| Bas - Pro.3 | Control | Bold | -0.36093048 | 0.7219419 | 33 | -0.49994391 | 1.000000000 |
| Bas - Pro.4 | Control | Bold | 0.45334310 | 0.7934421 | 33 | 0.57136258 | 1.000000000 |
| Pro.1 - Pro.2 | Control | Bold | -0.05945533 | 0.7305548 | 33 | -0.08138380 | 1.000000000 |
| Pro.1 - Pro.3 | Control | Bold | 0.02660331 | 0.7038053 | 33 | 0.03779925 | 1.000000000 |
| Pro.1 - Pro.4 | Control | Bold | 0.84087689 | 0.8022154 | 33 | 1.04819342 | 1.000000000 |
| Pro.2 - Pro.3 | Control | Bold | 0.08605864 | 0.7013562 | 33 | 0.12270318 | 1.000000000 |
| Pro.2 - Pro.4 | Control | Bold | 0.90033222 | 0.5963581 | 33 | 1.50971742 | 1.000000000 |
| Pro.3 - Pro.4 | Control | Bold | 0.81427358 | 0.7021554 | 33 | 1.15967708 | 1.000000000 |
| Bas - Pro.1 | Treatment | Bold | -0.62785360 | 0.7603440 | 33 | -0.82574941 | 1.000000000 |
| Bas - Pro.2 | Treatment | Bold | -0.50073160 | 0.7888211 | 33 | -0.63478472 | 1.000000000 |
| Bas - Pro.3 | Treatment | Bold | -0.31372887 | 0.7219419 | 33 | -0.43456246 | 1.000000000 |
| Bas - Pro.4 | Treatment | Bold | -0.89712636 | 0.7934421 | 33 | -1.13067660 | 1.000000000 |
| Pro.1 - Pro.2 | Treatment | Bold | 0.12712200 | 0.7305548 | 33 | 0.17400748 | 1.000000000 |
| Pro.1 - Pro.3 | Treatment | Bold | 0.31412474 | 0.7038053 | 33 | 0.44632334 | 1.000000000 |
| Pro.1 - Pro.4 | Treatment | Bold | -0.26927276 | 0.8022154 | 33 | -0.33566142 | 1.000000000 |
| Pro.2 - Pro.3 | Treatment | Bold | 0.18700273 | 0.7013562 | 33 | 0.26663018 | 1.000000000 |
| Pro.2 - Pro.4 | Treatment | Bold | -0.39639476 | 0.5963581 | 33 | -0.66469250 | 1.000000000 |
| Pro.3 - Pro.4 | Treatment | Bold | -0.58339749 | 0.7021554 | 33 | -0.83086658 | 1.000000000 |

| contrast | treatment | personality | estimate | SE | df | t.ratio | p.value |
|---|---|---|---|---|---|---|---|
| Bas - Pro.1 | Control | Shy | -0.17688236 | 0.9087849 | 33 | -0.19463612 | 1.000000000 |
| Bas - Pro.2 | Control | Shy | 0.22455110 | 0.9428216 | 33 | 0.23816924 | 1.000000000 |
| Bas - Pro.3 | Control | Shy | 0.88886000 | 0.8628857 | 33 | 1.03010171 | 1.000000000 |
| Bas - Pro.4 | Control | Shy | 1.00152355 | 0.9483446 | 33 | 1.05607551 | 1.000000000 |
| Pro.1 - Pro.2 | Control | Shy | 0.40143347 | 0.8731800 | 33 | 0.45973736 | 1.000000000 |
| Pro.1 - Pro.3 | Control | Shy | 1.06574236 | 0.8412083 | 33 | 1.26691858 | 1.000000000 |
| Pro.1 - Pro.4 | Control | Shy | 1.17840591 | 0.9588308 | 33 | 1.22900300 | 1.000000000 |
| Pro.2 - Pro.3 | Control | Shy | 0.66430889 | 0.8382810 | 33 | 0.79246561 | 1.000000000 |
| Pro.2 - Pro.4 | Control | Shy | 0.77697244 | 0.7127843 | 33 | 1.09005274 | 1.000000000 |
| Pro.3 - Pro.4 | Control | Shy | 0.11266355 | 0.8392363 | 33 | 0.13424533 | 1.000000000 |
| Bas - Pro.1 | Treatment | Shy | 2.57591450 | 0.7603440 | 33 | 3.38782780 | 0.018374438 |
| Bas - Pro.2 | Treatment | Shy | 2.82756698 | 0.7888211 | 33 | 3.58454770 | 0.010752783 |
| Bas - Pro.3 | Treatment | Shy | 3.05000546 | 0.7219419 | 33 | 4.22472401 | 0.001770278 |
| Bas - Pro.4 | Treatment | Shy | 2.78482636 | 0.7934421 | 33 | 3.50980435 | 0.013196619 |
| Pro.1 - Pro.2 | Treatment | Shy | 0.25165248 | 0.7305548 | 33 | 0.34446762 | 1.000000000 |
| Pro.1 - Pro.3 | Treatment | Shy | 0.47409096 | 0.7038053 | 33 | 0.67361094 | 1.000000000 |
| Pro.1 - Pro.4 | Treatment | Shy | 0.20891186 | 0.8022154 | 33 | 0.26041866 | 1.000000000 |
| Pro.2 - Pro.3 | Treatment | Shy | 0.22243848 | 0.7013562 | 33 | 0.31715479 | 1.000000000 |
| Pro.2 - Pro.4 | Treatment | Shy | -0.04274062 | 0.5963581 | 33 | -0.07166939 | 1.000000000 |
| Pro.3 - Pro.4 | Treatment | Shy | -0.26517910 | 0.7021554 | 33 | -0.37766438 | 1.000000000 |

Personality Variation in Associative Learning and Memory Retention in Zebrafish

(*Danio rerio*)

Princess Sunday-Jimmy[1]

Tran Thien An Le[1]

*Ryan Y. Wong[1],

[1]University of Nebraska at Omaha Biology Department, Omaha, NE USA

**Supplementary Table S4**: Raw behavioral measures recorded during the final 5 min of each conditioned place preference trial, including personality, sex, treatment, time spent, frequency of entry, and latency to first entry into the conditioned and non-conditioned zones.

| Fish ID | Trial | Sex | Group | Personality | Conditioned Stimulus for Time Spent (seconds) | Non-Conditioned Stimulus for Time Spent (seconds) | Conditioned Stimulus for Frequency of Entry | Non-Conditioned Stimulus for Frequency of Entry | Conditioned Stimulus for latency to First | Non-Conditioned Stimulus for latency to First |
|---|---|---|---|---|---|---|---|---|---|---|
| A_1 | Probe 1 | Male | Control | Bold | 157.567 | 134.233 | 10 | 10 | 3.233 | 0 |
| A_2 | Probe 1 | Male | Treatment | Bold | 90.2332 | 208.466 | 22 | 22 | 5.233 | 0 |
| A_3 | Probe 1 | Male | Treatment | Bold | 166.167 | 126.266 | 21 | 21 | 3.966 | 0 |
| A_4 | Probe 1 | Female | Control | Bold | 167.6 | 132.333 | 14 | 15 | 2.366 | 0 |
| A_5 | Probe 1 | Female | Treatment | Bold | 136.5 | 155.867 | 16 | 15 | 0 | 11.4 |
| A_1 | Baseline | Male | Control | Bold | 105.766 | 184.833 | 18 | 17 | 0 | 3.366 |
| A_2 | Baseline | Male | Treatment | Bold | 141.033 | 157.466 | 10 | 10 | 1.333 | 0 |
| A_3 | Baseline | Male | Treatment | Bold | 137.633 | 153.7 | 18 | 18 | 0 | 7.4 |
| A_4 | Baseline | Female | Control | Bold | 125.966 | 173.133 | 18 | 18 | 0 | 6.333 |
| A_5 | Baseline | Female | Treatment | Bold | 112.233 | 179.366 | 29 | 28 | 0 | 0.333 |
| A_1 | Probe 2 | Male | Control | Bold | 137.067 | 152.933 | 10 | 11 | 5.633 | 0 |
| A_2 | Probe 2 | Male | Treatment | Bold | 110.7 | 188.333 | 22 | 24 | 6.366 | 0 |
| A_3 | Probe 2 | Male | Treatment | Bold | 159.2 | 132.766 | 20 | 20 | 14.2 | 0 |
| A_4 | Probe 2 | Female | Control | Bold | 117.6 | 180.9 | 15 | 16 | 8.866 | 0 |
| A_5 | Probe 2 | Female | Treatment | Bold | 142.9 | 148.333 | 12 | 12 | 0 | 0.766 |
| A_1 | Probe 3 | Male | Control | Bold | 144.9 | 144.433 | 13 | 13 | 1.5 | 0 |
| A_2 | Probe 3 | Male | Treatment | Bold | 81.5666 | 212.933 | 9 | 9 | 10.733 | 0 |
| A_3 | Probe 3 | Male | Treatment | Bold | 181.966 | 110.2 | 15 | 15 | 0 | 0.6 |
| A_4 | Probe 3 | Female | Control | Bold | 59.9333 | 239.333 | 7 | 8 | 181.833 | 0 |
| A_5 | Probe 3 | Female | Treatment | Bold | 124.467 | 167.366 | 13 | 14 | 4.9 | 0 |
| A_1 | Probe 4 | Male | Control | Bold | 131.2 | 153.067 | 8 | 7 | 0 | 5.533 |
| A_2 | Probe 4 | Male | Treatment | Bold | 119.933 | 179.833 | 16 | 15 | 0 | 3.533 |

| | | | | | | | | | | |
|---|---|---|---|---|---|---|---|---|---|---|
| A_3 | Probe 4 | Male | Treatment | Bold | 195.366 | 103.466 | 11 | 11 | 7.7 | 0 |
| A_4 | Probe 4 | Female | Control | Bold | 190.933 | 99.3329 | 16 | 16 | 4.066 | 0 |
| A_5 | Probe 4 | Female | Treatment | Bold | 143.233 | 155.3 | 22 | 21 | 0 | 3.566 |
| B_12 | Baseline | Female | Control | Shy | 131.466 | 167.4 | 27 | 26 | 0 | 2.3 |
| B_17 | Baseline | Male | Treatment | Shy | 0 | 298.733 | 0 | 1 | 300 | 0 |
| B_18 | Baseline | Female | Treatment | Shy | 173.6 | 125.833 | 22 | 22 | 1.866 | 0 |
| B_19 | Baseline | Male | Treatment | Shy | 1.76666 | 298.23334 | 0 | 0 | 300 | 300 |
| B_20 | Baseline | Female | Treatment | Shy | 93.1332 | 206.299 | 10 | 11 | 88.8 | 0 |
| B_12 | Probe 1 | Female | Control | Shy | 177.966 | 121.633 | 13 | 13 | 5.033 | 0 |
| B_17 | Probe 1 | Male | Treatment | Shy | 244.799 | 55.201 | 1 | 0 | 0 | 300 |
| B_18 | Probe 1 | Female | Treatment | Shy | 172.466 | 126.6 | 18 | 18 | 0 | 0.7 |
| B_19 | Probe 1 | Male | Treatment | Shy | 1.3 | 298.7 | 1 | 1 | 0 | 0 |
| B_20 | Probe 1 | Female | Treatment | Shy | 106.266 | 192.966 | 12 | 11 | 0 | 2.433 |
| B_12 | Probe 2 | Female | Control | Shy | 207.033 | 91.1663 | 15 | 15 | 6.133 | 0 |
| B_17 | Probe 2 | Male | Treatment | Shy | 107.8 | 191.433 | 14 | 13 | 0 | 2.366 |
| B_18 | Probe 2 | Female | Treatment | Shy | 76.7666 | 222.166 | 20 | 21 | 32.733 | 0 |
| B_19 | Probe 2 | Male | Treatment | Shy | 149.066 | 150.067 | 25 | 25 | 0 | 2.733 |
| B_20 | Probe 2 | Female | Treatment | Shy | 208.166 | 91.2663 | 15 | 15 | 6.3 | 0 |
| B_12 | Probe 3 | Female | Control | Shy | 211.299 | 88.2332 | 19 | 18 | 0 | 3.033 |
| B_17 | Probe 3 | Male | Treatment | Shy | 236.499 | 56.6999 | 5 | 4 | 0 | 19.033 |
| B_18 | Probe 3 | Female | Treatment | Shy | 176.7 | 122.5 | 18 | 17 | 0 | 7.666 |
| B_19 | Probe 3 | Male | Treatment | Shy | 166.1 | 133.167 | 20 | 19 | 0 | 15.466 |
| B_20 | Probe 3 | Female | Treatment | Shy | 175.633 | 123.933 | 14 | 14 | 8.366 | 0 |
| B_12 | Probe 4 | Female | Control | Shy | 209.733 | 89.1666 | 14 | 14 | 0 | 41.733 |
| B_17 | Probe 4 | Male | Treatment | Shy | 211.266 | 87.9332 | 11 | 11 | 0 | 22.1 |
| B_18 | Probe 4 | Female | Treatment | Shy | 0 | 298.866 | 0 | 1 | 300 | 0 |
| B_19 | Probe 4 | Male | Treatment | Shy | 177.466 | 121.1 | 15 | 15 | 25.2 | 0 |
| B_20 | Probe 4 | Female | Treatment | Shy | 179.5 | 119.8 | 16 | 15 | 0 | 6.6 |
| C_1 | Baseline | Male | Control | Bold | 98.5996 | 200.766 | 16 | 16 | 0 | 1.433 |
| C_2 | Baseline | Female | Control | Bold | 249 | 49.433 | 23 | 24 | 10.4 | 0 |
| C_3 | Baseline | Male | Control | Bold | 160.133 | 138.233 | 13 | 13 | 18.766 | 0 |

| | | | | | | | | | | |
|---|---|---|---|---|---|---|---|---|---|---|
| C_4 | Baseline | Male | Control | Bold | 120.5 | 178.8 | 25 | 26 | 21.966 | 0.033 |
| C_5 | Baseline | Male | Treatment | Bold | 141.333 | 157.967 | 17 | 17 | 0 | 4.1 |
| C_1 | Probe 1 | Male | Control | Bold | 172.433 | 127.033 | 10 | 11 | 9.433 | 0 |
| C_2 | Probe 1 | Female | Control | Bold | 212.6 | 86.7329 | 18 | 18 | 1.133 | 0 |
| C_3 | Probe 1 | Male | Control | Bold | 137.833 | 160.333 | 14 | 14 | 24 | 0 |
| C_4 | Probe 1 | Male | Control | Bold | 48.7666 | 250.433 | 21 | 22 | 9.2 | 0 |
| C_5 | Probe 1 | Male | Treatment | Bold | 208.233 | 90.9666 | 16 | 16 | 0 | 14.5 |
| C_1 | Probe 2 | Male | Control | Bold | 76.9666 | 221.666 | 13 | 14 | 14.366 | 0 |
| C_2 | Probe 2 | Female | Control | Bold | 227.199 | 72.1333 | 17 | 16 | 0 | 24.766 |
| C_3 | Probe 2 | Male | Control | Bold | 200.1 | 97.3996 | 9 | 9 | 1.9 | 0 |
| C_4 | Probe 2 | Male | Control | Bold | 132.5 | 166.5 | 21 | 20 | 0 | 7.7 |
| C_5 | Probe 2 | Male | Treatment | Bold | 269.766 | 29.5 | 21 | 20 | 0 | 16.366 |
| C_1 | Probe 3 | Male | Control | Bold | 127.1 | 163.633 | 14 | 14 | 0 | 7 |
| C_2 | Probe 3 | Female | Control | Bold | 226.8 | 72.7996 | 14 | 15 | 0.666 | 0 |
| C_3 | Probe 3 | Male | Control | Bold | 297.3 | 0.466666 | 1 | 1 | 0 | 0 |
| C_4 | Probe 3 | Male | Control | Bold | 118.5 | 178.633 | 13 | 14 | 0.733 | 0 |
| C_5 | Probe 3 | Male | Treatment | Bold | 165.666 | 133.567 | 26 | 25 | 0 | 9.966 |
| C_1 | Probe 4 | Male | Control | Bold | 93.9666 | 201.333 | 16 | 17 | 2.7 | 0 |
| C_2 | Probe 4 | Female | Control | Bold | 196.933 | 102.3 | 15 | 15 | 0 | 34.7 |
| C_3 | Probe 4 | Male | Control | Bold | 164 | 135.266 | 13 | 14 | 4.833 | 0 |
| C_4 | Probe 4 | Male | Control | Bold | 134.233 | 164.667 | 18 | 17 | 0 | 21.6 |
| C_5 | Probe 4 | Male | Treatment | Bold | 197.366 | 101.467 | 18 | 17 | 0 | 4.833 |
| C_11 | Baseline | Male | Control | Shy | 128.2 | 167.2 | 19 | 19 | 0 | 21.4 |
| C_15 | Baseline | Female | Treatment | Shy | 112.2 | 187.566 | 18 | 17 | 0 | 1.4 |
| C_16 | Baseline | Male | Treatment | Shy | 153.8 | 146 | 6 | 6 | 0 | 103.966 |
| C_11 | Probe 1 | Male | Control | Shy | 106.7 | 191.8 | 9 | 10 | 21.566 | 0 |
| C_15 | Probe 1 | Female | Treatment | Shy | 45.3 | 253.833 | 5 | 5 | 227.666 | 0 |
| C_16 | Probe 1 | Male | Treatment | Shy | 299.466 | 0 | 1 | 0 | 0 | 300 |
| C_11 | Probe 2 | Male | Control | Shy | 1.4 | 294.366 | 1 | 2 | 28.566 | 0 |
| C_15 | Probe 2 | Female | Treatment | Shy | 168.033 | 131.1 | 16 | 16 | 0 | 4.933 |
| C_16 | Probe 2 | Male | Treatment | Shy | 214.1334 | 75.8666 | 24 | 23 | 0 | 15.5 |

| | | | | | | | | | | |
|---|---|---|---|---|---|---|---|---|---|---|
| C_11 | Probe 3 | Male | Control | Shy | 298.766 | 0 | 1 | 0 | 0 | 300 |
| C_15 | Probe 3 | Female | Treatment | Shy | 92.0999 | 207.233 | 19 | 19 | 6.566 | 0 |
| C_16 | Probe 3 | Male | Treatment | Shy | 278.899 | 19.9333 | 17 | 17 | 0 | 0.566 |
| C_11 | Probe 4 | Male | Control | Shy | 156.133 | 142.966 | 8 | 8 | 10.4 | 0 |
| C_15 | Probe 4 | Female | Treatment | Shy | 173.866 | 124.8 | 16 | 16 | 0 | 1.5 |
| C_16 | Probe 4 | Male | Treatment | Shy | 227.033 | 71.7333 | 19 | 18 | 0 | 2.966 |
| B_1 | Baseline | Female | Control | Bold | 123.9 | 174.966 | 32 | 32 | 2.933 | 0 |
| B_2 | Baseline | Male | Control | Bold | 123.333 | 175.866 | 15 | 15 | 0 | 5.2 |
| B_3 | Baseline | Female | Control | Bold | 192.033 | 106.833 | 32 | 32 | 0 | 1.533 |
| B_4 | Baseline | Male | Control | Bold | 106.966 | 191.9 | 12 | 12 | 0 | 0.5 |
| B_5 | Baseline | Male | Treatment | Bold | 54.1666 | 244.966 | 15 | 16 | 0.666 | 0 |
| B_6 | Baseline | Female | Treatment | Bold | 153.133 | 143.4 | 21 | 20 | 0 | 10.6 |
| B_8 | Baseline | Female | Treatment | Bold | 129.066 | 170.033 | 20 | 20 | 0 | 2.6 |
| B_10 | Baseline | Female | Treatment | Bold | 122.133 | 177.1 | 19 | 19 | 2.4 | 0 |
| B_13 | Baseline | Male | Control | Shy | 137.5 | 161.5 | 24 | 24 | 0 | 5.366 |
| B_14 | Baseline | Female | Control | Shy | 160.366 | 139.1 | 13 | 13 | 0 | 9.3 |
| B_15 | Baseline | Male | Control | Shy | 89.2999 | 209.633 | 7 | 8 | 116.433 | 0 |
| B_21 | Baseline | Male | Treatment | Shy | 3.833 | 289 | 0 | 0 | 300 | 300 |
| B_1 | Probe 1 | Female | Control | Bold | 97.9999 | 200.733 | 34 | 35 | 11.133 | 0 |
| B_2 | Probe 1 | Male | Control | Bold | 106.533 | 192.7 | 13 | 12 | 0 | 2.433 |
| B_3 | Probe 1 | Female | Control | Bold | 121.066 | 178.533 | 27 | 26 | 0 | 2.866 |
| B_4 | Probe 1 | Male | Control | Bold | 125.7 | 173.866 | 10 | 10 | 0 | 6.566 |
| B_5 | Probe 1 | Male | Treatment | Bold | 134.733 | 164.067 | 22 | 21 | 0 | 1.066 |
| B_6 | Probe 1 | Female | Treatment | Bold | 117.067 | 181.533 | 15 | 16 | 24.933 | 0 |
| B_8 | Probe 1 | Female | Treatment | Bold | 165.466 | 133.6 | 17 | 17 | 2 | 0 |
| B_10 | Probe 1 | Female | Treatment | Bold | 143.866 | 155.267 | 32 | 32 | 0 | 4.166 |
| B_13 | Probe 1 | Male | Control | Shy | 124.333 | 174.766 | 21 | 21 | 0 | 0.666 |
| B_14 | Probe 1 | Female | Control | Shy | 188.466 | 110.433 | 14 | 15 | 0.5 | 0 |
| B_15 | Probe 1 | Male | Control | Shy | 176.233 | 122.3 | 14 | 14 | 0 | 4.7 |
| B_21 | Probe 1 | Male | Treatment | Shy | 260.5663 | 29.6333 | 1 | 1 | 0 | 23.566 |
| B_1 | Probe 2 | Female | Control | Bold | 72.2666 | 226.999 | 29 | 30 | 0.633 | 0 |

| | | | | | | | | | | |
|---|---|---|---|---|---|---|---|---|---|---|
| B_2 | Probe 2 | Male | Control | Bold | 143.933 | 155.2 | 19 | 19 | 0 | 9.733 |
| B_3 | Probe 2 | Female | Control | Bold | 66.0666 | 233.533 | 17 | 17 | 12.333 | 0 |
| B_4 | Probe 2 | Male | Control | Bold | 136.833 | 162.033 | 14 | 14 | 27 | 0 |
| B_5 | Probe 2 | Male | Treatment | Bold | 174.7 | 123.833 | 17 | 16 | 0 | 0.3 |
| B_6 | Probe 2 | Female | Treatment | Bold | 283.466 | 16.0666 | 10 | 9 | 0 | 7.233 |
| B_8 | Probe 2 | Female | Treatment | Bold | 92.1332 | 206.733 | 27 | 28 | 4.6 | 0 |
| B_10 | Probe 2 | Female | Treatment | Bold | 143.2 | 156.433 | 21 | 21 | 14.333 | 0 |
| B_13 | Probe 2 | Male | Control | Shy | 139.466 | 160.133 | 21 | 20 | 0 | 2.633 |
| B_14 | Probe 2 | Female | Control | Shy | 148.233 | 151.367 | 14 | 14 | 0 | 9.233 |
| B_15 | Probe 2 | Male | Control | Shy | 201.899 | 96.6999 | 13 | 13 | 0 | 42.933 |
| B_21 | Probe 2 | Male | Treatment | Shy | 35.4 | 258.333 | 1 | 2 | 214.133 | 0 |
| B_1 | Probe 3 | Female | Control | Bold | 55.9666 | 242.566 | 18 | 19 | 9.833 | 0 |
| B_2 | Probe 3 | Male | Control | Bold | 96.6332 | 202.366 | 17 | 17 | 3.9 | 0 |
| B_3 | Probe 3 | Female | Control | Bold | 82.5999 | 216.233 | 18 | 18 | 9.733 | 0 |
| B_4 | Probe 3 | Male | Control | Bold | 159.466 | 139.9 | 9 | 9 | 0 | 11.366 |
| B_5 | Probe 3 | Male | Treatment | Bold | 178.2 | 121 | 16 | 16 | 0 | 24.566 |
| B_6 | Probe 3 | Female | Treatment | Bold | 261.066 | 38.6333 | 7 | 6 | 0 | 34.766 |
| B_8 | Probe 3 | Female | Treatment | Bold | 50.233 | 248.933 | 37 | 37 | 0 | 2.5 |
| B_10 | Probe 3 | Female | Treatment | Bold | 191.366 | 108.233 | 27 | 27 | 0 | 10.733 |
| B_13 | Probe 3 | Male | Control | Shy | 148.5 | 150 | 17 | 18 | 7.866 | 0 |
| B_14 | Probe 3 | Female | Control | Shy | 172.933 | 126.566 | 16 | 16 | 2.366 | 0 |
| B_15 | Probe 3 | Male | Control | Shy | 130.467 | 169.3 | 10 | 11 | 15.866 | 0 |
| B_21 | Probe 3 | Male | Treatment | Shy | 162.133 | 136.3 | 19 | 19 | 0 | 1.333 |
| B_1 | Probe 4 | Female | Control | Bold | 113.466 | 185.933 | 21 | 20 | 0 | 3.666 |
| B_2 | Probe 4 | Male | Control | Bold | 142.233 | 157.2 | 20 | 21 | 1 | 0 |
| B_3 | Probe 4 | Female | Control | Bold | 296.8333 | 1.4 | 2 | 1 | 0 | 0 |
| B_4 | Probe 4 | Male | Control | Bold | 175.5 | 123.533 | 10 | 10 | 0 | 7.666 |
| B_5 | Probe 4 | Male | Treatment | Bold | 188.833 | 107.7 | 12 | 11 | 0 | 5.9 |
| B_6 | Probe 4 | Female | Treatment | Bold | 288.566 | 11.0333 | 6 | 5 | 0 | 39.6 |
| B_8 | Probe 4 | Female | Treatment | Bold | 58.2999 | 240.899 | 31 | 32 | 54.533 | 0 |
| B_10 | Probe 4 | Female | Treatment | Bold | 54.4999 | 244.833 | 6 | 7 | 188.4 | 0 |

| | | | | | | | | | | |
|---|---|---|---|---|---|---|---|---|---|---|
| B_13 | Probe 4 | Male | Control | Shy | 131.867 | 167.6 | 12 | 12 | 8.566 | 0 |
| B_14 | Probe 4 | Female | Control | Shy | 158.833 | 138.533 | 13 | 12 | 0 | 5.4 |
| B_15 | Probe 4 | Male | Control | Shy | 115.433 | 184.166 | 6 | 6 | 0 | 16.566 |
| B_21 | Probe 4 | Male | Treatment | Shy | 41.06666 | 257.39999 | 2 | 2 | 0 | 0 |
| C_6 | Baseline | Male | Treatment | Bold | 61.9666 | 237.299 | 31 | 32 | 3.866 | 0 |
| C_7 | Baseline | Male | Treatment | Bold | 116.766 | 182.233 | 24 | 23 | 0 | 12.633 |
| C_12 | Baseline | Male | Control | Shy | 24.4333 | 267.433 | 3 | 4 | 219.933 | 0 |
| C_14 | Baseline | Male | Control | Shy | 157.433 | 142.033 | 14 | 14 | 39.066 | 0 |
| C_18 | Baseline | Male | Treatment | Shy | 0.6 | 294.1 | 0 | 1 | 300 | 0 |
| C_19 | Baseline | Male | Treatment | Shy | 0 | 297.833 | 0 | 1 | 300 | 0 |
| C_20 | Baseline | Male | Treatment | Shy | 0.3 | 298.599 | 1 | 2 | 157.733 | 0 |
| C_6 | Probe 1 | Male | Treatment | Bold | 32.5666 | 266.399 | 15 | 16 | 13.233 | 0 |
| C_7 | Probe 1 | Male | Treatment | Bold | 146.166 | 153.567 | 19 | 19 | 0 | 0.966 |
| C_12 | Probe 1 | Male | Control | Shy | 131.067 | 168.2 | 15 | 15 | 57.333 | 0 |
| C_14 | Probe 1 | Male | Control | Shy | 0 | 298.833 | 0 | 1 | 300 | 0 |
| C_18 | Probe 1 | Male | Treatment | Shy | 3 | 287.233 | 0 | 1 | 300 | 0 |
| C_19 | Probe 1 | Male | Treatment | Shy | 119.4 | 179.433 | 13 | 13 | 0 | 7.8 |
| C_20 | Probe 1 | Male | Treatment | Shy | 129.1 | 168.466 | 17 | 17 | 8.533 | 0 |
| C_6 | Probe 2 | Male | Treatment | Bold | 70.8996 | 228.433 | 23 | 23 | 0 | 0.933 |
| C_7 | Probe 2 | Male | Treatment | Bold | 167.533 | 131.4 | 16 | 15 | 0 | 7.233 |
| C_12 | Probe 2 | Male | Control | Shy | 157 | 142.766 | 9 | 10 | 6.766 | 0 |
| C_14 | Probe 2 | Male | Control | Shy | 79.0999 | 219.533 | 10 | 11 | 130.8 | 0 |
| C_18 | Probe 2 | Male | Treatment | Shy | 178.666 | 120.133 | 23 | 22 | 0 | 6.866 |
| C_19 | Probe 2 | Male | Treatment | Shy | 155.866 | 75.8666 | 24 | 23 | 0 | 15.5 |
| C_20 | Probe 2 | Male | Treatment | Shy | 186.433 | 112.867 | 19 | 18 | 0 | 1.833 |
| C_6 | Probe 3 | Male | Treatment | Bold | 166.533 | 132.7 | 21 | 22 | 13.6 | 0 |
| C_7 | Probe 3 | Male | Treatment | Bold | 178.233 | 120.733 | 14 | 14 | 0.466 | 0 |
| C_12 | Probe 3 | Male | Control | Shy | 69.5996 | 229.333 | 7 | 7 | 0 | 18.733 |
| C_14 | Probe 3 | Male | Control | Shy | 163.433 | 135.033 | 18 | 18 | 3.1 | 0 |
| C_18 | Probe 3 | Male | Treatment | Shy | 278.899 | 19.9333 | 17 | 17 | 0 | 0.566 |
| C_19 | Probe 3 | Male | Treatment | Shy | 145.867 | 152.966 | 20 | 21 | 3.3 | 0 |

| | | | | | | | | | | |
|---|---|---|---|---|---|---|---|---|---|---|
| C_20 | Probe 3 | Male | Treatment | Shy | 142.167 | 156.8 | 21 | 21 | 17.8 | 0 |
| C_6 | Probe 4 | Male | Treatment | Bold | 101.9 | 197 | 16 | 17 | 4.533 | 0 |
| C_7 | Probe 4 | Male | Treatment | Bold | 169.633 | 130.2 | 14 | 14 | 0 | 5.133 |
| C_12 | Probe 4 | Male | Control | Shy | 140.267 | 159.5 | 18 | 18 | 7.6 | 0 |
| C_14 | Probe 4 | Male | Control | Shy | 131.7 | 167.566 | 15 | 15 | 0 | 7.933 |
| C_18 | Probe 4 | Male | Treatment | Shy | 185.8 | 113.267 | 8 | 7 | 0 | 9.8 |
| C_19 | Probe 4 | Male | Treatment | Shy | 170.833 | 128.2 | 12 | 13 | 9.3 | 0 |
| C_20 | Probe 4 | Male | Treatment | Shy | 122.333 | 176.9 | 13 | 12 | 0 | 2.933 |
